# T cells exhibit unexpectedly low discriminatory power and can respond to ultra-low affinity peptide-MHC ligands

**DOI:** 10.1101/2020.11.14.382630

**Authors:** Johannes Pettmann, Enas Abu-Shah, Mikhail Kutuzov, Daniel B. Wilson, Michael L. Dustin, Simon J. Davis, P. Anton van der Merwe, Omer Dushek

## Abstract

T cells use their T cell receptors (TCRs) to discriminate between peptide MHC (pMHC) ligands that bind with different affinities but precisely how different remains controversial. This is partly because the affinities of physiologically relevant interactions are often too weak to measure. Here, we introduce a surface plasmon resonance protocol to measure ultra-low TCR/pMHC affinities (K_D_ ~ 1000 *μ*M). Using naïve, memory, and blasted human CD8^+^ T cells we find that their discrimination power is unexpectedly low, in that they require a large >100-fold decrease in affinity to abolish responses. Interestingly, the discrimination power reduces further when antigen is presented in isolation on artificial surfaces but can be partially restored by adding ligands to CD2 or LFA-1. We were able to fit the kinetic proof-reading model to our data, yielding the first estimates for both the time delay (2.8 s) and number of biochemical steps (2.67). The fractional number of steps suggest that one of the proof-reading steps is not easily reversible.

## Introduction

T cells use their T cell receptors (TCRs) to recognise short peptides presented on Major Histocompatibility Complex (pMHC). The ability of T cells to discriminate between self and foreign derived pMHCs is the cornerstone of adaptive immunity and defects in this process can lead to autoimmunity or immunodeficiency. Therefore, the discrimination power of T cells and its underlying mechanisms have been extensively studied but remain a topic of debate (1–3).

Previous studies using three murine TCRs (OT-I, 2B4 and 3.L2) have suggested that T cells exhibit remarkable discrimination powers (4–8). It was reported that mutations to the cognate peptide that produced only modest 3–7-fold decreases in affinity completely abolished the T cell response even when the peptide concentration was increased by as much as 100,000-fold on the antigen-presenting cell (APC). This could also be observed using plate-immobilised recombinant pMHC (in the absence of APCs) (5), suggesting that discrimination can proceed by TCR/pMHC interactions without other co-signalling receptors. This striking ‘perfect’ discrimination cannot be explained by occupancy models, where a linear relationship is predicted between the dissociation constant (K_D_) and pMHC potency (i.e. a 3-fold increase in K_D_ can be compensated for by a 3-fold increase in pMHC concentration). Although the standard kinetic proofreading (KP) mechanism can explain the nonlinear discrimination (9, 10), it failed to also explain the high sensitivity of T cells that enables them to respond to 1–10 pMHC ligands (11–15). This is due to a trade-off between discrimination and sensitivity, so that parameters that explain high sensitivity cannot explain high levels of discrimination (16). This prompted work focused on augmenting KP with additional mechanisms to explain how high levels of discrimination can be achieved while retaining sensitivity (17–30).

Although not used for the quantitative study of antigen discrimination, a large number of other TCRs have now been characterised. These other TCRs appear to exhibit a more graded loss of T cell responses as the affinity decreases (e.g. 1G4 TCR (31), B3K506 TCR (32), and 1E6 TCR (33); see also Huhn et al (34)). Therefore, there is an apparent discrepancy in the strength of discrimination between studies. Moreover, discrimination has not been assessed in primary quiescent human T cells.

A key challenge in assessing discrimination is the accurate measurements of TCR/pMHC affinities, which like other surface interactions on leukocytes can be very weak, with K_D_ ranging from 1 to >100 *μ*M (35). A highly sensitive and widely-used method for analysing molecular interactions is surface plasmon resonance (SPR) but even with this method, accurate measurements are difficult to make, especially at 37°C. In the case of 2B4 and 3.L2, measurements were performed at 25°C where off-rates are slower and affinities typically higher. However, non-linear changes with temperatures have been documented so that fold-changes between two interactions can differ between 25°C and 37°C (31). In the case of OT-1, measurements were performed at 37°C but high affinity bi-phasic binding was observed (6), which has not been observed for other TCRs and may represent protein aggregates that often form at the high concentrations necessary for making these measurements. It follows that the reported 3-fold change in affinity between the activating OVA and non-activating E1 pMHC (6) may be a consequence of multivalent interactions. Indeed, more recent studies found the expected low-affinity mono-phasic binding for OT-1/OVA (36, 37) and no detectable binding for OT-1/E1 (36). These studies highlight the challenges of accurately measuring TCR/pMHC affinities and underline their importance in understanding antigen discrimination.

In this work we introduce a new SPR protocol that can accurately determine ultra-low TCR/pMHC affinities at 37°C into the K_D_ ~ 1000 *μ*M regime. Using panels of pMHCs for two different human TCRs, we found that discrimination is surprisingly low with responses detected even to ultra-low affinity antigens. By directly fitting the KP model to all data, we find that it can simultaneously explain sensitivity and discrimination with a fast proofreading time delay and a small, fractional number of biochemical steps. The work reconciles discrepancies between previous studies and the surprisingly low discrimination power has implications for both basic and translational T cell immunology.

## Results

### Measurements of ultra-low TCR/pMHC affinities at 37°C

To quantitatively assess antigen discrimination, we first generated ligands to the anti-tumour 1G4 (38) and anti-viral A6 (39) TCRs recognising peptides loaded on HLA-A*02:01 and measured affinities using SPR at 37°C. The standard SPR protocol is based on injecting the TCR at increasing concentrations over a surface with immobilised pMHC (Figure 1A,B). The steady-state binding response at each concentration is then plotted over the TCR concentration (Figure 1C). This curve is fitted by a 2-parameter Hill function to determine B_max_ (the maximum response when all pMHC are bound by TCR) and the K_D_, which is the TCR concentration where the binding response is half the B_max_. Therefore, accurate determination of K_D_ requires an accurate determination of B_max_.

**Figure 1:**
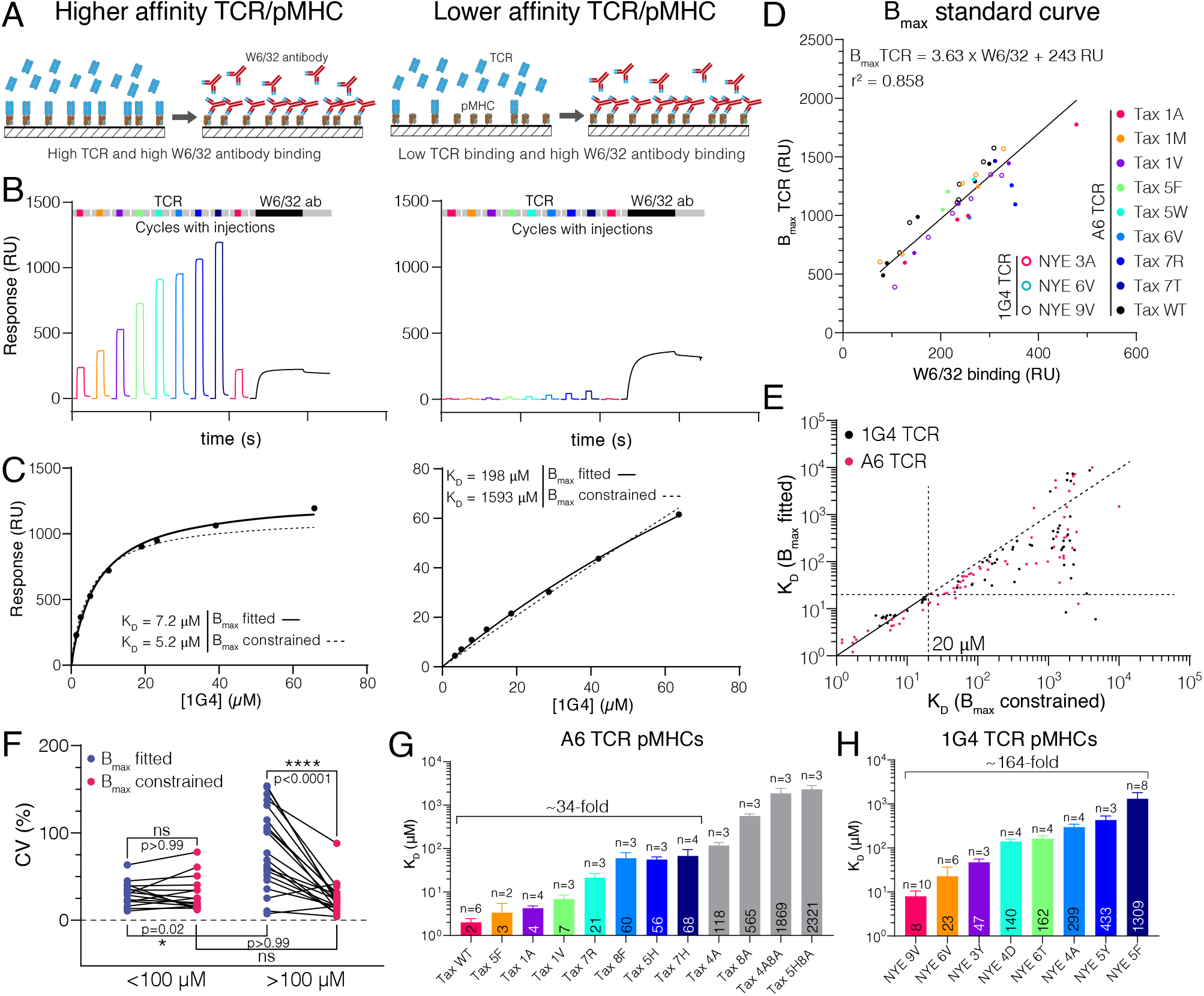
Measuring ultra-low TCR/pMHC affinities using SPR at 37°C using a constrained B_max_ method. **(A-C)** Comparison of 1G4 TCR binding to a higher (left panels, 9V) and lower (right panels, 5F) affinity pMHC. **(A)** Schematic comparing binding of TCR and W6/32 to high and low affinity pMHC ligands. **(B)** Example SPR sensograms showing binding to immobilised pMHC with sequential injections of different concentrations of TCR followed by the W6/32 antibody. **(C)** A plot of the steady-state binding response from (B) over the TCR concentration (filled circles) is fitted to determine K_D_ when B_max_ is either unconstrained (standard method) or constrained (new method). **(D)** Empirical standard curve relating the W6/32 binding response to fitted B_max_ values obtained using only higher affinity interactions. **(E)** Correlation of K_D_s obtained obtained using unconstrained (fitted) versus constrained B_max_ values. Each dot represents an individual measurement (n=136; 63 for 1G4 TCR, 73 for A6 TCR). **(F)** Coefficient of variation for higher (<100 *μ*M) or lower affinity (>100 *μ*M) interactions. Data was compared using a Kruskal-Wallis test with Dunn’s multiple comparison. **(G)** Selected pMHC panel for A6 TCR (mean with SD). **(H)** Selected pMHC panel for 1G4 TCR (mean with SD). Values of K_D_ are indicated in bars and ligands used for functional experiments in the main text are coloured.

In the case of the 1G4 TCR binding to its cognate NY-ESO-1 peptide, this protocol produces binding response curves that saturate, allowing for an accurate estimate of B_max_ and therefore an accurate estimate of K_D_ = 7.2 *μ*M (Figure 1A-C, left column), which agrees with previous reports (31). However, when the experiment is repeated for pMHCs that bind with lower affinity, the binding response curves do not saturate (Figure 1A-C, right column). Because of this the fitted B_max_ and therefore the fitted K_D_ may not be accurate. Previously, saturation was achieved by lowering the temperature (where affinities can be higher) and/or by using higher TCR concentrations, but both methods can produce artefacts (see Introduction).

To determine B_max_ for low affinity pMHCs, we generated a standard curve by taking advantage of the conformation-sensitive, pan-HLA-A/B/C antibody (W6/32) that only binds correctly folded pMHC (40, 41). By injecting saturating concentrations of W6/32 at the end of each experiment (Figure 1B, black line) we were able to plot the fitted B_max_ from higher affinity interactions (where binding saturated) over the maximum W6/32 binding (Figure 1D). We observed a linear relationship even when including data for different TCRs binding different pMHC across multiple protein preparations immobilised at different levels, strongly supporting the notion that the W6/32 antibody and the TCR recognise the same correctly-folded pMHC population. This justified the use of the standard curve based on W6/32 binding to estimate B_max_ for low affinity TCR interactions. We next fitted K_D_ values for 136 interactions using the standard method where B_max_ is fitted and the new method where B_max_ is constrained to the value obtained using the standard curve (Figure 1E). In the new method, the only fitted parameter is K_D_. Both methods produced similar K_D_ values for higher affinity interactions, validating the method. In contrast, large (100-fold) discrepancies appeared for lower affinity interactions, with the fitted B_max_ method consistently underestimating the K_D_ compared to the constrained B_max_ method (see Figure 1E). These large discrepancies were observed despite both methods providing a similar fit (e.g. Figure 1C, right). This means that for the fitted B_max_ method, different combinations of B_max_ and K_D_ can provide a fit of similar quality so that the fitted K_D_ can exhibit large variations for the same interaction (also known as ‘over-fitting’). We explored this by comparing the precision of both methods using the coefficient of variance (CV) of multiple measurements of the same TCR/pMHC combination. We found a similar CV for higher affinity interactions (<100 *μ*M K_D_) but the CV increased for lower affinity interactions when the B_max_ was fitted but not when the B_max_ was constrained (Figure 1F).

Taken together, this analysis shows that unlike the standard method, the new constrained B_max_ method allows reproducible K_D_ measurements even at low affinities. Moreover, it highlights the ability of SPR to measure ultra-low affinity interactions (K_D_ ~ 1000 *μ*M). This enabled us to identify panels of pMHCs for the A6 and 1G4 TCRs that spanned a very wide-range of affinities to use in antigen discrimination assays (Figure 1G,H).

### The discrimination power of T cells is surprisingly low

To quantify the degree of discrimination, we introduced the 1G4 TCR into quiescent naïve or memory CD8^+^ T cells that were then co-cultured with autologous monocyte-derived dendritic cells (moDCs) pulsed with each peptide (Figure 2A). Using surface CD69 as a marker for T cell activation, we found that lowering the affinity gradually reduced the response without the sharp affinity threshold predicted by perfect discrimination, and remarkably, responses could be measured to ultra-low affinity peptides, such as NYE 5F (K_D_ ~ 1309 *μ*M; see Figure 2B,C). To exclude that this may be a result of preferential loading and/or stability of ultra-low affinity peptides, we pulsed the TAP-deficient T2 cell lines with all peptides and found similar HLA upregulation, suggesting similar peptide presentation (Figure S1). We quantified the functional potency as the concentration of peptide required to reach 15 % activation (P_15_), which correlated with K_D_ (Figure 2D,E).

**Figure 2:**
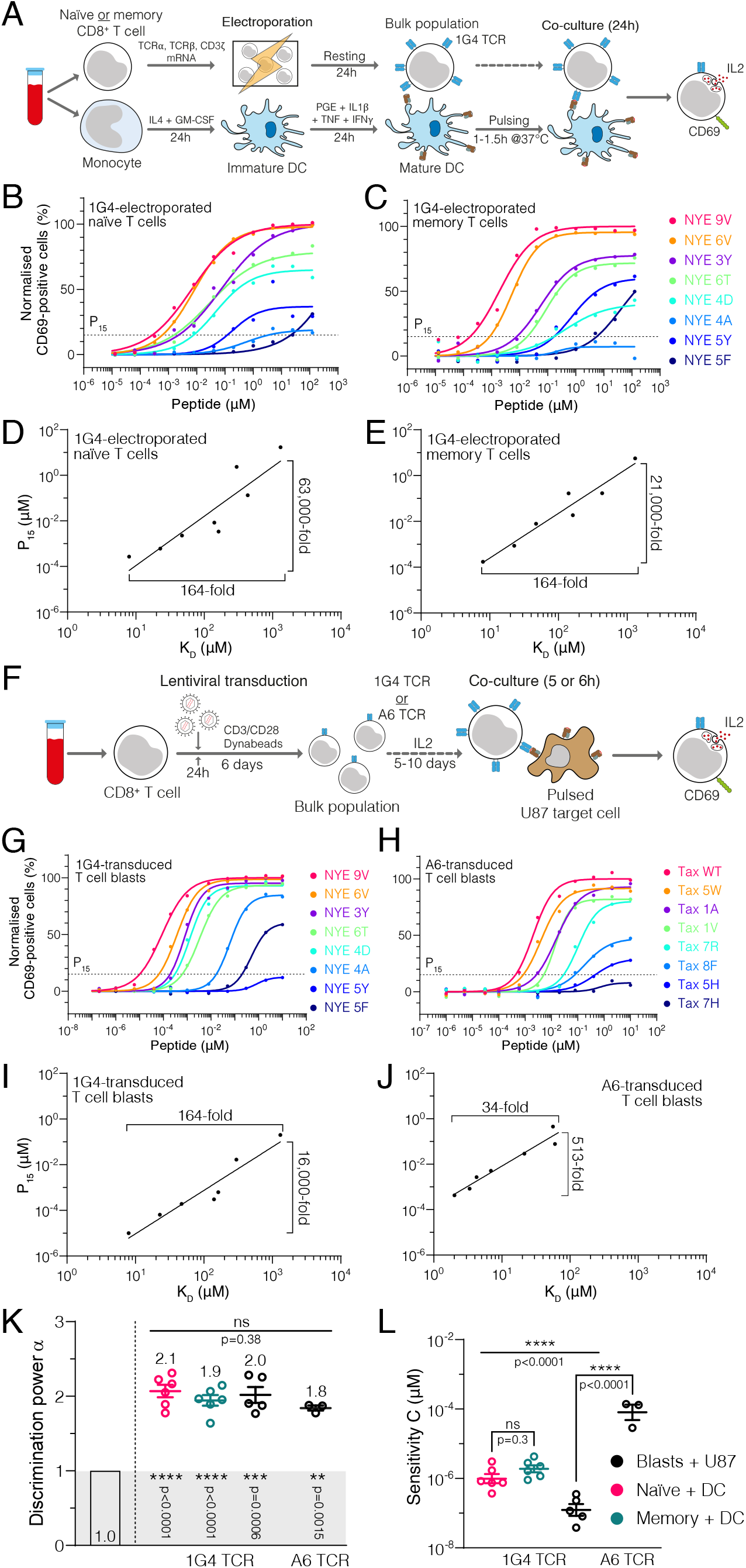
Naïve, memory, and blast human CD8^+^ T cells exhibit surprisingly low discrimination powers. **(A)** Experimental protocol for studying quiescent primary human naïve and memory CD8^+^ T cells interacting with autologous moDCs as APCs. **(B,C)** Example dose-responses for naïve and memory T cells with sigmoidal fit. Potency (P15) is determined by the concentration of peptide eliciting 15 % activation (dotted line). **(D,E)** Examples of potency vs. K_D_ fitted with a power law for naïve and memory T cells. Fold-change in K_D_ and in potency derived from fits are shown. **(F)** Experimental protocol for studying primary human CD8^+^ T cell blasts interacting with the glioblastoma cell line U87 as APCs. Example dose-responses **(G,H)** and potency vs. K_D_ plots **(I,J)** for T cell blasts expressing the indicated 1G4 or A6 TCR. **(K-L)** Comparison of the fitted discrimination power (α) and fitted sensitivity (*C*). Shown is the mean with SEM with each dot representing an individual experiment (n=3–6). In **(K)**, conditions were compared with a 1-way ANOVA and each condition was compared to α=1 (baseline discrimination provided by an occupancy model) with an independent 1-sample Student’s t test. In **(L)**, the 1G4 data was compared with a ordinary 1-way ANOVA and all data was compared using a second ordinary 1-way ANOVA with Sidak’s multiple comparison for a pairwise test.

We next investigated discrimination in CD8^+^ T cell blasts Figure 2F, which serve as an *in vitro* model for effector T cells and are commonly used in adoptive cell therapy. 1G4-expressing T cell blasts also exhibited a graded loss of the response and, as before, activation could be observed to ultra-low affinity peptides (Figure 2G). To independently corroborate antigen discrimination with a second TCR, we repeated these experiments with A6-expressing T cell blasts, and again found a graded response (Figure 2H). Unlike 1G4 T cell blasts, responses were only observed for higher affinity peptides with K_D_s <100 *μ*M (Figure 2H, Figure S2A,B). We attribute this to the much lower expression of the A6 TCR (Figure S2C,D). Nonetheless, potency correlated with affinity (Figure 2I,J).

We next used the potency/affinity relationships to quantify discrimination. We first noted that some level of discrimination can be observed because the ~160-fold variation in affinity for the 1G4 TCR was amplified into a 16,000–63,000-fold variation in potency (Figure 2D,E,I). To quantify the non-linearity, we fit the potency plots with a power law,

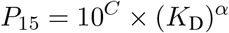

where α measures the discrimination power (slope on the log-log plot) and *C* measures sensitivity (y-intercept on the log-log plot). The baseline discrimination power provided by an occupancy model is α = 1 and analysis of the original mouse TCR data produces α ~ 9 (34). Although higher than baseline, we found surprisingly low discrimination powers (1.8–2.1), that were similar for naïve, memory, and blasted T cells and for both the 1G4 and A6 TCRs (Figure 2K). Interestingly, the sensitivities varied considerably, as evident in the ~1,000-fold lower sensitivity for the A6 T cell blasts (Figure 2L). In addition to CD69 expression, we examined the production of IL-2 as a measure of T cell activation and found similar discrimination powers (Figure S3A-C). In summary, we found that the discriminatory power of the TCR is lower than previously reported and appears to be independent of the expression level of the TCR, the maturation status of the T cell, and the sensitivity to antigen.

### The discrimination power of T cells is dependent on ligands for co-signalling receptors

We next sought to determine if antigen discrimination is influenced by engagement of co-signalling receptors. Since APCs express ligands for many co-signalling receptors, we adopted a reductionist system (42, 43) where recombinant pMHCs are presented on plates with or without ligands for co-signalling receptors before being used as stimulation surfaces for 1G4 T cell blasts (Figure 3A).

**Figure 3:**
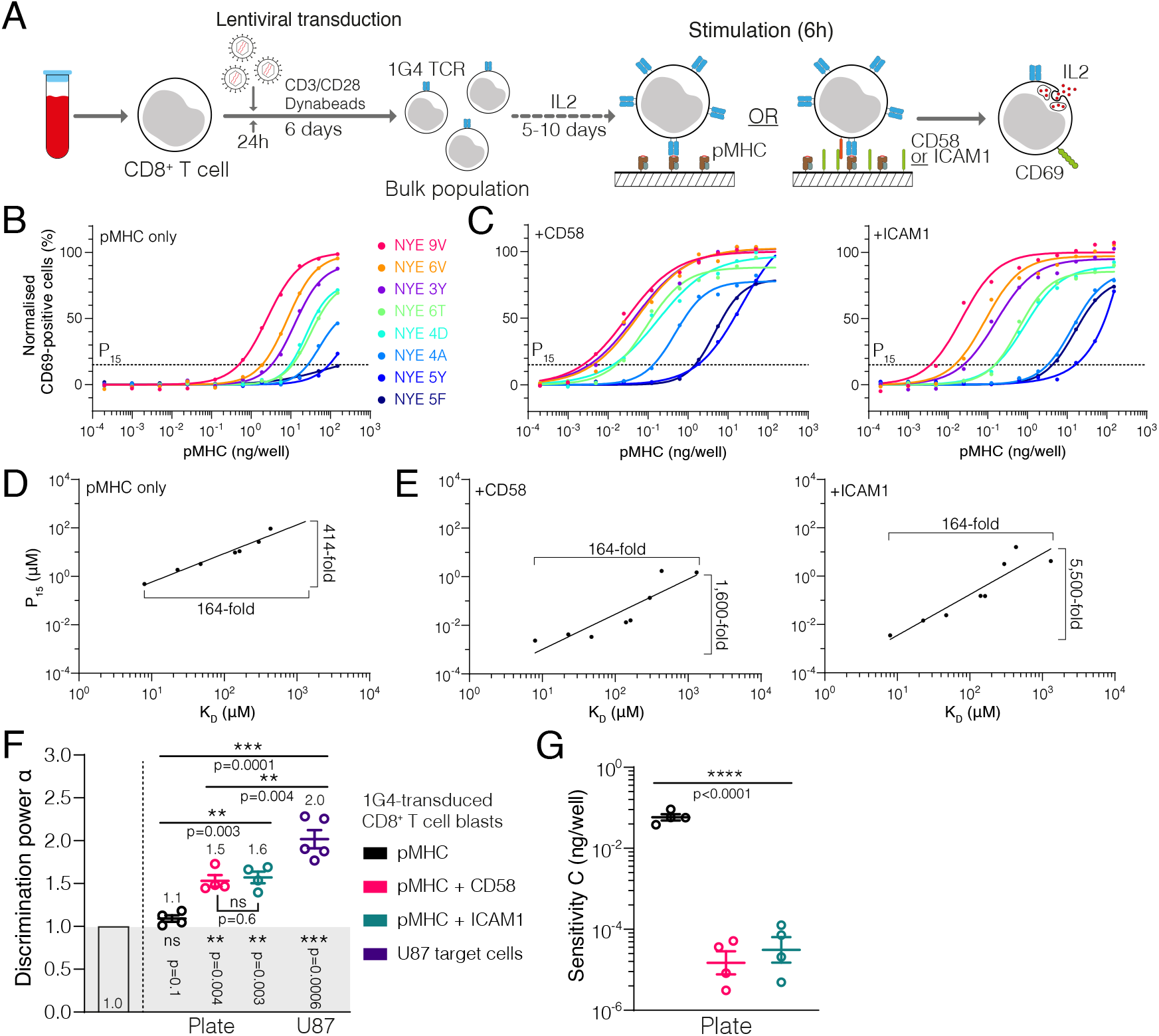
The discrimination power of T cells is enhanced by ligation of the receptors CD2 or LFA-1. **(A)** Experimental protocol for stimulation of primary human CD8^+^ T cell blasts with plate-bound recombinant ligands. **(B,C)** Example dose-response curve for 1G4 T cell blasts stimulated with (B) pMHC alone or (C) in combination with CD58 or ICAM1. **(D,E)** Potency derived from dose-response curves over K_D_ showing the power function fit (D) with pMHC alone or (E) in combination with CD58 or ICAM1. **(F)** Comparison of discrimination power α (mean with SEM, n=4–5). U87 data as in Figure 2K. Plate data was compared using repeated-measure 1-way ANOVA (Geisser-Greenhouse corrected) with Sidak’s comparison for indicated pairwise comparisons. CD58 and ICAM1 were compared to U87 co-culture data using ordinary 1-way ANOVA. U87 and pMHC alone were compared using a Student’s t-test. Each condition was compared to the discrimination power of an occupancy model (α=1) using each an independent 1-sample Student’s t test. **(G)** Comparison of sensitivity *C* (mean with SEM, n=4–5). Comparison using repeated-measure 1-way ANOVA (Geisser-Greenhouse corrected).

As before, the T cell response was gradually reduced with lower affinity pMHCs and responses could be detected to ultra-low affinity pMHCs (Figure 3B). A plot of the potency over the affinity revealed that a 164-fold variation in affinity led only to a 414-fold variation in potency (Figure 3D), which was substantially lower than the 16,000-fold variation observed for T cell blasts with APCs (e.g. Figure 2I). This significant reduction in the TCR discrimination power when pMHC is presented in isolation compared to APCs is quantified by the fitted α decreasing from 2.0 to 1.1 (Figure 3F).

We hypothesised that the absence of ligands for co-signalling receptors may account for this reduction in antigen discrimination. It has been shown that ligation of co-signalling receptors can greatly enhance antigen potency (42, 44). Inclusion of recombinant ICAM1 (a ligand of LFA-1) or CD58 (the ligand to CD2) increased TCR downregulation (Figure S4) and antigen potency (Figure 3C) in this experimental system, consistent with previous reports using APCs (44, 45). As before, we fit the power relationship to the potency vs. affinity plots (Figure 3E), which allowed us to make two striking quantitative observations. First, ligation of either receptor increased discrimination from 414-fold variation in potency to >1600-fold (Figure 3D vs. Figure 3E). This is reflected in the discrimination power, which increased from 1.1 to 1.5 and 1.6 (Figure 3F). Second, ligation of either receptor increased antigen potency (or sensitivity) by 100-fold, which is appreciably larger than previous reports (44) and likely reflects the reductionist system we have used where other co-signalling receptors cannot compensate. These two observations were reproduced when T cell activation was assessed by IL-2 secretion (Figure S3).

Taken together, these results reveal that the discrimination power of the TCR reduces to baseline when recognising antigen in isolation and that the co-signalling receptors CD2 and LFA-1 enhance not only antigen sensitivity but also antigen discrimination.

### Kinetic proofreading explains the discrimination power with a fast proofreading time delay and a fractional number of steps

KP is an operational model proposed to explain the discrimination power of T cells (9). It relies on a sequence of biochemical steps that introduce a time delay (*τ*_KP_) between the initial binding step (step 0) and the final signalling step (step N). In addition, it relies on pMHC unbinding instantly reversing all steps returning the TCR to its basal state (Figure 4A). Despite being introduced more than 20 years ago and underlying all models of T cell activation (46), there are no estimates of either the number of steps or the time delay for T cells discriminating antigens on APCs. These parameters not only provide molecular insights but also determine whether the KP mechanism is sufficient to explain discrimination (16, 19, 23, 26).

**Figure 4:**
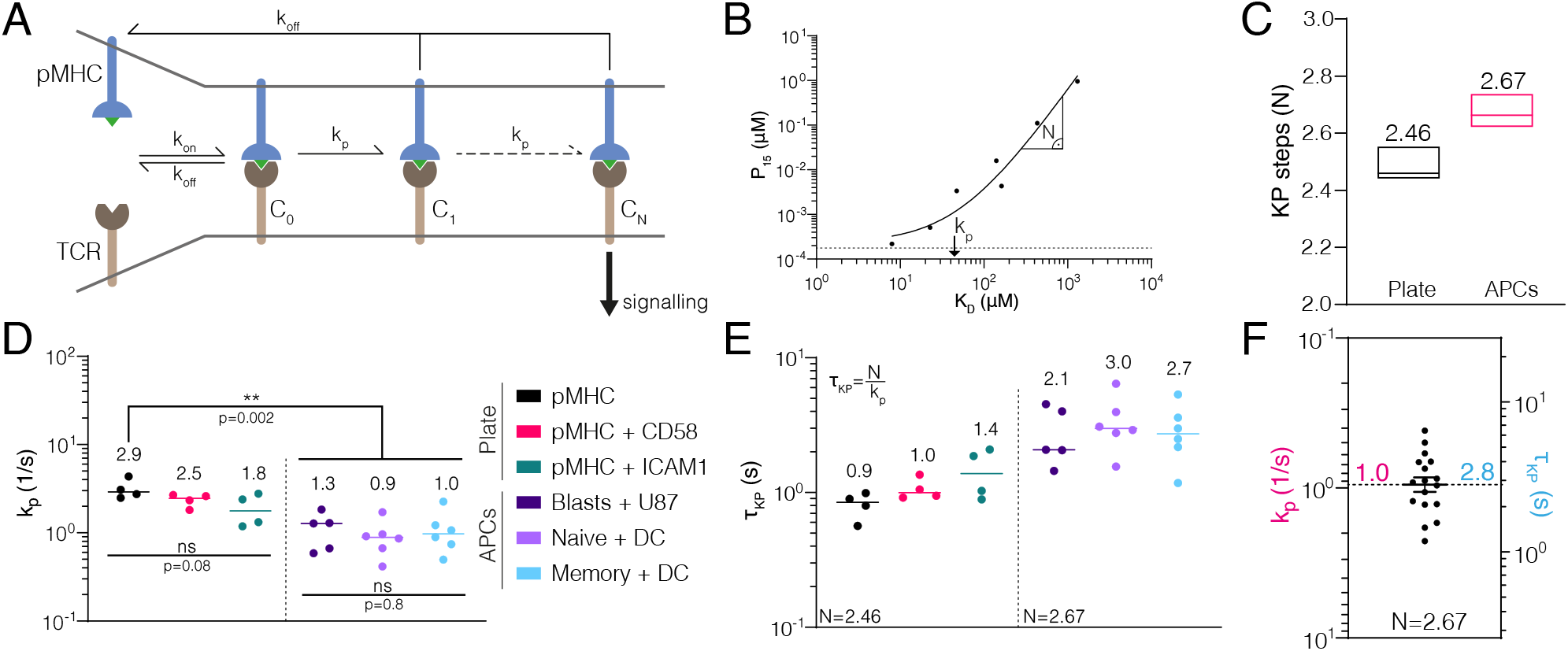
Direct fit of kinetic proofreading reveals fractional steps with a short proofreading time delay. **(A)** The kinetic proofreading (KP) model showing that upon TCR binding to pMHC (step 0) a sequence of N steps with rate *k_p_* are performed before productive signalling (step N) with all steps immediately reversed upon pMHC unbinding (with rate *k*_off_). The pMHC binding time required to produce a TCR signal is the kinetic proofreading time delay defined as *τ_KP_* = *N* /*k_p_*. **(B)** Example fit of the KP model showing that the fitted *k_p_* is approximately the inflection point and *N* is approximately the slope away from the inflection point. **(C)** The fitted number of steps was a global shared parameter for all plate data or all APC data. The median value of N with the min/max produced by the global fit is shown. **(D)** The proofreading rate was a local parameter for individual data sets. Each data point is the median value produced by the fit to each data set. Plate data compared using a repeated-measure 1-way ANOVA (Geisser-Greenhouse corrected) and APC data and APC vs. plate data was compared using each an ordinary 1-way ANOVA. **(E)** The proofreading time was calculated using the fitted value of N in panel C and the individual values of *k_p_* in panel D. **(F)** Pooled APC data (naïve, memory, blast T cells) are used to compute the mean value of *k_p_* and *τ*_KP_. Error bars show SEM.

In order to estimate these parameters, we directly fitted the KP model to either all the APC data or all the plate data separately. The KP model parameters included the number of steps (*N*) and the rate of each step (*k_p_*), which together determine the proofreading time delay (*τ*_KP_ = *N* /*k_p_*), and can be visualised on the potency plots (Figure 4B). The KP model produced an excellent fit to each data set (e.g. Figure 4B, Figure S6A, B) and, importantly, the fit method we implemented showed that *N* and *k_p_* can be uniquely determined (Figure S6C-H).

We found an unexpectedly small and fractional number of biochemical steps when fitting the APC data (2.67), and a similar value when independently fitting the plate data (Figure 4C). A similar number of steps (2.7 ± 0.5) was recently reported for KP performed by a chimeric antigen receptor (CAR) (47). Although a small number of steps is surprising given that the TCR is known to undergo a large number of biochemical modifications, it can be explained by the fact that only steps that must be performed sequentially contribute to the proofreading chain (see Discussion).

However, it was not intuitive how a fractional number of biochemical steps can arise. We explored the possibility that this can arise if at least one intermediate step was not instantly reversible (termed step *K* here). In this model, the modification in step K is sustained for a period of time after pMHC unbinds so that a population of TCR can exist with some proofreading steps already achieved. It follows that pMHC binding to this population of TCR would short-circuit KP undergoing only *N* - *K* steps before reaching the final step. The apparent number of steps would then be a weighted average between N and *N* - *K*, depending on the relative amount of TCR with K steps sustained, so that fractions are possible (e.g. with *N* = 3 and *K* = 1, we can arrive at effectively 2.5 steps). We solved this modified KP model and found that it can transition between 3 and 2 steps (Figure S7). Therefore, a fractional number of steps can suggest that one (or more) KP steps may not be instantly reversible and instead can be sustained upon pMHC unbinding.

We found that the fitted *k_p_* values were similar within the APC experiments but generally slower than the plate experiments (Figure 4D), and because a similar number of steps were observed in both, this observation translated to the time delay which was longer on the APCs (Figure 4E). Therefore, the higher discrimination power observed on APCs compared to the plate (Figure 3F) is a result of a longer time delay (9, 16) produced not by more steps but rather a slower rate of each step. This makes conceptual sense because the number of steps is constrained by the signalling architecture whereas the rate of each step can be regulated. We found that all APC data produced similar KP parameters, and therefore combined these values to provide an average time delay of *τ*_KP_ = 2.8 s using *N* = 2.67 (Figure 4F).

We next sought to determine whether these KP parameters, which we have shown can explain the discrimination power, can also explain antigen sensitivity. To do this, we used a previously described measure of sensitivity ([*k_p_*/(*k_p_* + *k*_off_)]^*N*^) (16), which is the fraction of times that pMHC binding reaches the final signalling state. Using *k*_off_ = 0.33 s^−1^ (1G4 binding to 9V (31)) and the averaged KP parameters above, we find that 45 % of pMHC binding events will reach the final productive signalling state consistent with high sensitivity.

## Discussion

Using a new SPR protocol, we accurately measured ultra-low TCR/pMHC affinities finding T cell responses to these antigens at ultra-high concentrations. This enabled us to measure the discriminatory power of TCR recognition with unprecedented accuracy. Quantifying the discrimination power produces values of 1.9–2.1 using APCs. In a separate analysis of the published literature, we found similar discrimination powers for other human and mouse TCRs (2.0 with 95% CI of 1.5-2.4) that were significantly lower than those for the original murine TCR data (~9.0) (34). The discrimination power we report is significantly higher than the baseline predicted by a receptor occupancy model (1.0) and can readily be explained by a KP mechanism.

The discrepancy between previous work (4–8, 19, 25, 27) and other human and mouse TCRs (34), including the present work, is likely a result of inaccurate SPR affinity measurements. In addition to temperature and protein aggregation (see Introduction), we have found that extrapolating the K_D_ from non-saturating binding curves reduces precision and underestimates the K_D_. Given that fitting programs report extrapolated K_D_ values without warnings, it is important that caution is used when interpreting these. By constraining B_max_ using a standard curve based on the W6/32 antibody binding, we avoided this, enabling us to provide accurate K_D_ values into the ultra-low 1000 *μ*M regime. This method may be useful to measure TCR binding to low-affinity self pMHCs.

We have found that the KP mechanism can simultaneously explain both high sensitivity and the discrimination power of T cells. This is achieved by a few fractional steps (2.67) and a short proofreading time delay (2.8 s). This time delay is at the lower end of the range reported using soluble tetramers (8 s with 95 % CI: 3–19 s) (48) and consistent with the 4 s time delay between pMHC binding and LAT phosphorylation (49). The small number of steps is reasonable because, although the TCR undergoes a large number of biochemical modifications (2, 3), only steps that must be sequential contribute. It follows that individual ITAMs acting in parallel cannot extend the proofreading chain. This is consistent with the fact that the number of steps we report for the TCR (10 ITAMs) is the same as the number reported for a CAR (6 ITAMs) (47). Moreover, the fact that this number is fractional suggests that one (or more) step(s) is not easily reversible. This step may be ZAP70 recruitment, which has a lifetime at the membrane that is longer than the TCR/pMHC lifetime (50–52) and therefore, it may sustain intermediate signalling for a period of time. This ability is consistent with reports suggesting that the TCR can integrate signals across multiple rebinding events (21, 25, 31, 32).

The KP mechanism couples discrimination and sensitivity so that increases in sensitivity lead to decreases in discrimination (16). Unexpectedly, we found similar discrimination powers yet large differences in sensitivity (up to 1000-fold) between T cell states (e.g. naïve/memory vs. blast), between two TCRs (1G4 vs. A6), and between readouts (CD69 vs. IL-2). This suggests that the coupling between discrimination and sensitivity may be lower than previously thought. Consistent with this, we found that ligation of CD2 or LFA-1 dramatically increased sensitivity and increased the discrimination power. These observations can be reconciled with the KP mechanism if sensitivity is not limited by the proofreading time delay but rather by downstream signalling modules that set the effective threshold on the number of productive TCR signals required to activate T cells. Put differently, T cells may operate in a discrimination regime that produces TCR signals even with low affinity ligands but the decision to respond is made further downstream, and can be regulated both by the T cell state and by engagement of co-signalling receptors.

The level of antigen discrimination we report in this study and in an analysis of the literature (34) can be explained by the standard KP model. A number of studies, including work from our lab, have reported additional mechanisms beyond the standard KP model to explain the higher level of discrimination originally reported (17–32). Given that these mechanisms augment the standard proofreading model, they can all explain the lower level of discrimination that we report. Future work is needed to dissect the precise contribution of each molecular mechanism to the overall discrimination power of T cells.

The broad range of mechanisms that can produce high discrimination powers (17–32) suggests the observed low discrimination power may be a feature rather than a design flaw of T cells. A possible feature is the ability to perform expression-level discrimination of self pMHCs so that T cells can recognise and appropriately respond to cells over-expressing self antigens. Recently, a compelling theory proposed that expression-level discrimination is required to maintain homeostasis of endocrine tissues by killing hyper-secreting mutants (53). The flip-side of a lower discrimination power is the critical requirement for additional regulatory mechanisms to maintain tolerance, such as the established mechanisms of co-signalling receptors and T_regs_ (54). The finding that co-signalling receptors can control not only sensitivity but also discrimination, suggests new avenues to optimise T cell discrimination for therapeutic purposes.

## Materials and Methods

### Protein production

Class I pMHCs were refolded as previously described (55). Human HLA-A*0201 heavy chain (UniProt residues 25–298) with a C-terminal AviTag/BirA recognition sequence and human beta-2 microgolublin were expressed in *Escherichia coli* and isolated from inclusion bodies. Trimer was refolded by consecutively adding peptide, *β*2M and heavy chain into refolding buffer and incubating for 2–3 days at 4°C. Protein was filtered, concentrated using centrifugal filters, biotinylated (BirA biotin-protein ligase bulk reaction kit [Avidity, USA]) and purified by size exclusion chromatography (Superdex 75 column [GE Healthcare]) in HBS-EP (0.01 M HEPES pH 7.4, 0.15 M NaCl, 3 mM EDTA, 0.005 % v/v Tween20). Purified protein was aliquoted and stored at −80°C until use. Soluble α and β subunits of 1G4 and A6 TCRs were produced in *E. coli*, isolated from inclusion bodies, refolded *in vitro* and purified using size exclusion chromatography in HBS-EP, as described previously (31).

Soluble extracellular domain (ECD) of human CD58 (UniProt residues 29–204 or 29–213) was produced either in Freestyle 293F suspension cells (Thermo Fisher) or adherent, stable GS CHO cell lines. For the latter, cells were expanded in selection medium (10 % dialysed FCS, 1x GSEM supplement [Sigma-Aldrich], 20–50 *μ*M MSX, 1 % Pen/Strep) for at least 1 week. Production was performed in production medium (2–5 % FCS, 1x GSEM supplement, 20 *μ*M MSX, 2 mM sodium butyrate, 1 % Pen/Strep) continuously for a few weeks with regular medium exchanges. Production in 293F was performed according to manufacturer’s instructions. Human ICAM1 ECD (UniProt residues 28–480) was either produced by transient transfection or lentiviral transduction of adherent 293T, or by transient expression in 293F, as described above. All supernatants were 0.45 *μ*m filtered and 100 *μ*M PMSF was added. Proteins were purified using standard Ni-NTA agarose columns, followed by *in vitro* biotinylation as described above. Alternatively, ligands were biotinylated by co-transfection (1:10) of a secreted BirA-encoding plasmid and adding 100 *μ*M D-biotin to the medium, as described before (56). Proteins were further purified and excess biotin removed from proteins biotinylated *in vitro* by size exclusion chromatography (Superdex 75 or 200 column [GE Healthcare]) in HBS-EP; purified proteins were aliquoted and stored at −80°C until use.

Biotinylation levels of pMHC and accessory ligands were routinely tested by gel shift on SDS-PAGE upon addition of saturating amounts of streptavidin.

### SPR

TCR–pMHC interactions were analysed on a Biacore T200 instrument (GE Healthcare Life Sciences) at 37°C and a flow rate of 10 *μ*l/min. Running buffer was HBS-EP. Streptavidin was coupled to CM5 sensor chips using an amino coupling kit (GE Healthcare Life Sciences) to near saturation, typically 10000–12000 response units (RU). Biotinylated pMHCs were injected into the experimental flow cells (FCs) for different lengths of time to produce desired immobilisation levels (typically 500–1500 RU), which were matched as closely as feasible in each chip. Usually, FC1 was as a reference for FC2–FC4. Biotinylated CD58 ECD was immobilised in FC1 at a level matching those of pMHCs. In some experiments, another FC was used as a reference. Excess streptavidin was blocked with two 40 s injections of 250 *μ*M biotin (Avidity). Before TCR injections, the chip surface was conditioned with 8 injections of the running buffer. Dilution series of TCRs were injected simultaneously in all FCs; the duration of injections (3070 s) was the same for conditioning and TCR injections. Buffer was also injected after every 2 or 3 TCR injections; all binding data were double referenced (57) vs. the average of the closest buffer injections before and after TCR injection. After TCR injections, W6/32 antibody (10 *μ*g/ml; Biolegend) was injected for 10 min. Maximal W6/32 binding was used to generate the empirical standard curve and to infer the B_max_ of TCRs from the standard curve. The empirical standard curve only contained data where the ratio of the highest concentration of TCR to the fitted K_D_ value (obtained using the standard method with B_max_ fitted) was 2.5 or more. This threshold ensured that the binding response curves saturated so that only accurate measurements of B_max_ were included. All interactions were fit using both the fitted and constrained B_max_ method (Figure 1E). For constrained K_D_s above 20 *μ*M we reported the constrained K_D_, otherwise we use the B_max_ fitted K_D_.

### Co-culture of naïve & memory T cells

The assay was performed as previously described (58). Naïve and memory T cells were isolated from anonymized HLA-A2^+^ leukocyte cones obtained from the NHS Blood and Transplantation service at Oxford University Hospitals by (REC 11/H0711/7), using EasySep Human naïve CD8^+^ T Cell Isolation Kit (Stemcell) and EasySep Human Memory CD8^+^ T Cell Enrichment Kit (Stemcell), respectively. Cells were washed 3x with Opti-MEM serum-free medium (Thermo Fisher) and 2.5–5.0 Mio cells were resuspended at a density of 25 Mio/ml. Suspension was mixed with 5 *μ*g/Mio cells of 1G4α, 1G4β, and CD3ζ each and 100–200 *μ*l suspension was transferred into a BTX Cuvette Plus electroporation cuvette (2 mm gap; Harvard Bioscience). Electroporation was performed using a BTX ECM 830 Square Wave Electroporation System (Harvard Bioscience) at 300 V, 2 ms. T cells were used 24 h after electroporation.

Autologous monocytes were enriched from the same blood product using RosetteSep Human Monocyte Enrichment Cocktail (Stemcell), cultured at 1–2 Mio/ml in 12-well plates in the presence of 50 ng/ml IL4 (PeproTech) and 100 ng/ml GM-CSF (Immunotools) for 24 h to induce differentiation. Maturation into moDCs was induced by adding 1 *μ*M PGE_2_ (Sigma Aldrich), 10 ng/ml IL1β (Biotechne), 20 ng/ml IFNγ, and 50 ng/ml TNF (PeproTech) for an additional 24 h. MoDCs (50,000/well) were loaded for 60–90 min at 37°C with peptide and labelled with Cell Trace Violet (Thermo Fisher) to distinguish them from T cells prior to co-culturing with 50,000 T cells/well in a 96 well plate for 24 h. T cell activation was assessed by flow cytometry and testing culture supernatant for cytokines using ELISAs.

### T cell blasts

All cell culture of human T cells was done using complete RPMI (10 % FCS, 1 % penicillin/streptomycin) at 37°C, 5 % CO2. T cells were isolated from whole blood from healthy donors or leukocyte cones purchased from the NHS Blood and Transplantation service at the John Radcliffe Hospital. For whole blood donations, a maximum of 50 ml was collected by a trained phlebotomist after informed consent had been given. This project has been approved by the Medical Sciences Inter-Divisional Research Ethics Committee of the University of Oxford (R51997/RE001) and all samples were anonymised in compliance with the Data Protection Act.

For plate stimulations and experiments with U87 target cells, CD8^+^ T cells were isolated using RosetteSep Human CD8^+^ enrichment cocktail (Stemcell) at 6 *μ*l/ml for whole blood or 150 *μ*l/ml for leukocyte cones. After 20 min incubation at room temperature, blood cone samples were diluted 3.125-fold with PBS, while whole blood samples were used directly. Samples were layered on Ficoll Paque Plus (GE) at a 0.8:1.0 ficoll:sample ratio and spun at 1200 g for 20–30 min at room temperature. Buffy coats were collected, washed twice, counted and cells were resuspended in complete RMPI with 50 U/ml IL2 (PeproTech) and CD3/CD28 Human T-activator dynabeads (Thermo Fisher) at a 1:1 bead:cell ratio. Aliquots of 1 Mio cells in 1 ml medium were grown overnight in 12-or 24-well plates (either TC-treated or coated with 5 *μ*g/cm^2^ retronectin [Takara Bio]) and then transduced with VSV-pseudotyped lentivirus encoding for either the 1G4 or the A6 TCR. After 2 days (4 days after transduction), 1 ml of medium was exchanged, and IL2 was added to a final concentration of 50 U/ml. Beads were magnetically removed at day 5 post-transduction and T cells from thereon were resuspended at 1 Mio/ml with 50 U/ml IL2 every other day. For functional experiments, T cells were used between 10–16 days after transduction.

### Lentivirus production

HEK 293T or Lenti-X 293T (Takara) were seeded in complete DMEM in 6-well plate to reach 60–80 % confluency after one day. Cells were either transfected with 0.95 *μ*g pRSV-Rev, 0.37 *μ*g pVSV-G (pMD2.G), 0.95 *μ*g pGAG (pMDLg/pRRE), and 0.8 *μ*g of a EF1α promoter-transfer plasmid with 9 *μ*l X-tremeGENE 9 or HP (both Roche). Lentiviral supernatant was harvested after 20–30 h and filtered through a 0.45 *μ*m cellulose acetate filter. In an updated version, LentiX cells were transfected with 0.25 *μ*g pRSV-Rev, 0.53 *μ*g pGAG, 0.35 *μ*g pVSV-G, and 0.8 *μ*g transfer plasmid using 5.8 *μ*l X-tremeGENE HP. Medium was replaced after 12–18 h and supernatant harvested as above after 30–40 h. Supernatant from one well of a 6-well plate was used to transduce 1 Mio T cells.

### Co-culture of T cell blasts

For co-culture experiments with U87 (a kind gift of Vincenzo Cerundolo, University of Oxford), 30,000 target cells were seeded in a TC-coated 96-well F-bottom plate and incubated overnight. Peptides were diluted in complete DMEM (10 % FCS, 1 % penicillin/streptomycin) to their final concentration and incubated with U87 cells for 1–2 h at 37°C. Peptide-containing medium was removed and 60,000 TCR-transduced primary human CD8^+^ T cell blasts were added, spun for 2 min at 50 g, and incubated for 5 h at 37°C. At the end of the experiment, 10 mM EDTA was added and cells were detached by vigorous pipetting. Cells were stained for flow cytometry and analysed immediately, or fixed and stored for up to 1 day before running. Supernatants were saved for cytokine ELISAs.

### Plate stimulation

Glass-bottom Sensoplates (96-well; Greiner) were washed with 1 M HCl/70 % EtOH, thoroughly rinsed twice with PBS and coated overnight at 4°C with 100 *μ*l/well of 1 mg/ml biotinylated BSA (Thermo Fisher) in PBS. Plates were washed with PBS twice and incubated for at least 1 h with 20 *μ*g/ml streptavidin (Thermo Fisher) in 1 % BSA/PBS at room temperature. Plates were washed again with PBS and biotinylated pMHC (in-house) was added for at least 1 h at room temperature or overnight at 4°C. Plates were emptied and accessory ligand (CD58 or ICAM1, in-house) or PBS was added for the same duration as above. Upon completion, plates were washed once and stored for up to 1 day in PBS at 4°C.

For stimulation, T cells were counted, washed once to remove excess IL2, and 75,000 cells in 180–200 *μ*l complete RMPI were dispensed per well. Cells were briefly spun down at 50 g to settle to the bottom and subsequently incubated for 4 h at 37°C. At the end of the experiment, 10 mM EDTA was added and cells were detached by vigorous pipetting. Cells were stained for flow cytometry and analysed immediately, or fixed and stored for up to 1 day. Supernatants were saved for cytokine ELISAs.

### Peptides and loading

We used peptide ligands that were either described previously (31, 59–64) or designed by us based on the published crystal structures of these TCRs in complex with MHC (1G4: PDB 2BNQ, A6: PDB 1AO7). Peptides were synthesised at a purity of >95 % (Peptide Protein Research, UK). Tax WT is a 9 amino acid, class I peptide derived from HTLV-1 Tax_11–19_ (39, 65). NYE 9V refers to a heteroclitic (improved stability on MHC), 9 amino acid, class I peptide derived from the wild type NYE-ESO_157–165_ 9C peptide (38). See Table S1 and Table S2 for a list of peptides.

Loading efficiency was evaluated by pulsing T2 cells for 1–2 h at 37°C with a titration of peptides. Loading was assessed as upregulation of HLA-A2 (clone: BB7.2; Biolegend) by flow cytometry.

### Flow cytometry

Tetramers were produced in-house using refolded monomeric, biotinylated pMHC and streptavidin-PE (Biolegend) at a 1:4 molar ratio. Streptavidin-PE was added in 10 steps and incubated for 10 min while shaking at room temperature. Insoluble proteins were removed by brief centrifugation at 13,000 *g* and 0.05–0.1 % sodium azide added for preservation. Tetramers were kept for up to 3 months at 4°C. Cells were stained for CD69 with clones FN50 (Biolegend). Staining for CD45 (clone HI30; Biolegend) was used to distinguish target and effector cells in co-culture assays with U87 cells. Cell viability staining was routinely performed for plate stimulations and U87 co-culture using fixable violet or near-infrared viability dyes (Zombie UV fixable viability kit [Biolegend], Zombie NIR fixable viability kit [Biolegend], eBioscience fixable viability dye eFluor 780 [Invitrogen]). Samples were analysed using a BD X-20 flow cytometer and data analysis was performed using FlowJo v10 (BD Biosciences).

### ELISAs

Human IL-2 Ready-SET Go! ELISA kit (eBioscience/Invitrogen) or Human TNF alpha ELISA Ready-SET-Go! (eBioscience/Invitrogen) and Nunc MaxiSorp 96-well plates (Thermo Fisher) were used according to the manufacturer’s instructions to test appropriately diluted (commonly 4–30-fold) T cell supernatant for secretion of IL2 or TNF.

### TCR expression

TCRαβ-KO Jurkat E6.1 cells (a kind gift of Edward Jenkins) were transduced with 1G4 or A6 lentivirus and TCR expression was measured by staining for CD3 (clone: UCHT1; Biolegend) and TCRαβ (clone IP26; Biolegend).

### Data analysis & fitting

Details of statistical methods for comparisons can be found in the figure legends. All tests were 2-tailed, if not stated otherwise. All fitting was done using least squares optimisation, if not stated otherwise.

Quantitative analysis of antigen discrimination was performed by first fitting dose-response data with a 4-parameter sigmoidal model on a linear scale in Python v3.7 and lmfit v0.9.13 using Levenberg–Marquardt:

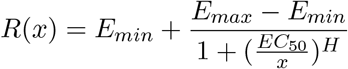

where *x* refers to the peptide concentration used to pulse the target cells (in *μ*M) or the amount of pMHC used to coat the well of a plate (in ng/well). The curve produced by this fit was used to interpolate potency as the concentration of antigen required to induce activation of 15 % for CD69 (*P*_15_) and 10 % for IL2 (*P*_10_). These percentages were chosen based on noise levels and to include lower affinity antigens in the potency plots. Potency values exceeding doses used for pulsing or coating were excluded from the analysis (i.e. no extrapolated data was included in the analysis).

To determine the discrimination power *α*, we fitted the power law in log-space to our data:

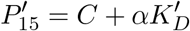

where 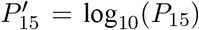 and 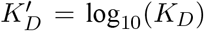. All data analysis was performed using GraphPad Prism 8 (GraphPad Software), if not stated otherwise.

For KP model fitting, we derived the following equation to relate potency to the model parameters:

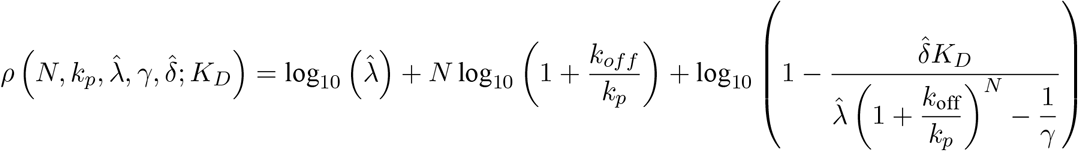

using custom C++ code (Apple LLVM version 7.0.0, clang-700.1.76). We fitted 2 independent datasets derived with either immobilised ligands, or APC co-culture. Parameters *N*, 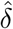 and *γ* were shared within each dataset. *k_p_* and 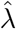 were fitted locally. See Supplementary Text for details.

This fitting procedure required an estimate of *k*_off_ for each peptide ligand but the fast kinetics of ultra-low affinity ligands meant that we were unable to resolve these using SPR. Instead, we noted that previous work has shown that on-rates exhibit small variations between pMHCs that differ by few amino acids (31, 59). We therefore estimated off-rates for each peptide ligand using the same on-rate (i.e. 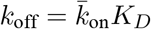) determined previously for 1G4 pMHCs (Figure S5).

## Acknowledgements

We thank Philipp Kruger, Edward Jenkins, Marcus Bridge, Samuel A. Isaacson, and Marion H. Brown for experimental and mathematical assistance.

## Contributions

JP: Conceptualisation, investigation, methodology, visualisation, formal analysis, data curation, project admin-istration, funding acquisition, writing – original draft, writing – review & editing. EAS: Conceptualisation, investigation, methodology, data curation, writing – review & editing, funding acquisition. MK: Conceptualisa-tion, investigation, resources, methodology, data curation, writing – review & editing. DBW: Investigation, data curation, software, formal analysis, writing – review & editing. SJD: Supervision, writing – review & editing, funding acquisition. MLD: Funding acquisition, supervision, writing – review & editing. PAvdM: Conceptuali-sation, funding acquisition, writing – review & editing. OD: Conceptualisation, project administration, funding acquisition, supervision, writing – original draft, writing – review & editing.

## Supplementary Information

**Table S1:** Fitted K_D_ with the indicated method for the 1G4 TCR in SPR at 37°C. Includes all peptides used for SPR standard curve and functional experiments.

| Name | Sequence | $K_D$ (Bmax fitted) | | | $K_D$ (Bmax const.) | | | n |
| --- | --- | --- | --- | --- | --- | --- | --- | --- |
| | | Mean (μM) | SD (μM) | CV ( %) | Mean (μM) | SD (μM) | CV ( %) | |
| NYE 9V | SLLMWITQV | 7.97 | 2.55 | 32.0 | 4.93 | 1.22 | 24.8 | 7 |
| NYE 3A | SLAMWITQV | 7.04 | 1.49 | 21.1 | 6.28 | 2.16 | 34.3 | 5 |
| NYE 6V | SLLMWVTQV | 17.6 | 5.28 | 30.0 | 22.8 | 13.8 | 60.8 | 6 |
| NYE 3Y | SLYMWITQV | 36.3 | 10.4 | 28.7 | 47.3 | 8.58 | 18.1 | 3 |
| NYE 4D | SLLDWITQV | 90.8 | 41.7 | 45.9 | 140 | 18.3 | 13.0 | 4 |
| NYE 6T | SLLMWTTQV | 84.6 | 30.3 | 35.9 | 162 | 26.3 | 16.2 | 4 |
| NYE 4A | SLLAWITQV | 157 | 80.8 | 51.4 | 299 | 48.6 | 16.3 | 4 |
| NYE 5Y | SLLMYITQV | 191 | 107 | 56.3 | 433 | 103 | 23.8 | 3 |
| NYE 5F | SLLMFITQV | 756 | 1092 | 144.5 | 1309 | 505 | 38.6 | 8 |

**Table S2:** Fitted K_D_ with the indicated method for the A6 TCR in SPR at 37°C. Includes all peptides used for SPR standard curve and functional experiments. N/A: not applicable.

| Name | Sequence | $K_D$ (Bmax fitted) | | | $K_D$ (Bmax const.) | | | n |
| --- | --- | --- | --- | --- | --- | --- | --- | --- |
| | | Mean (μM) | SD (μM) | CV ( %) | Mean (μM) | SD (μM) | CV ( %) | |
| Tax WT | LLFGYPVYV | 2.01 | 0.437 | 21.8 | 1.66 | 0.270 | 16.3 | 5 |
| Tax 5F | LLFGFPVYV | 3.36 | 2.12 | 63.2 | 2.67 | 2.08 | 77.9 | 2 |
| Tax 7T | LLFGYPTYV | 3.42 | N/A | N/A | 2.80 | N/A | N/A | 1 |
| Tax 5W | LLFGWPVYV | 6.08 | N/A | N/A | 5.00 | N/A | N/A | 1 |
| Tax 1M | MLFGYPVYV | 6.11 | N/A | N/A | 6.10 | N/A | N/A | 1 |
| Tax 1A | ALFGYPVYV | 4.25 | 0.571 | 13.4 | 6.50 | 0.902 | 13.9 | 4 |
| Tax 1V | VLFGYPVYV | 6.92 | 1.63 | 23.5 | 7.97 | 1.62 | 20.3 | 3 |
| Tax 6V | LLFGYVVYV | 13.5 | N/A | N/A | 21.1 | N/A | N/A | 1 |
| Tax 7R | LLFGYPRYV | 11.5 | 1.85 | 16.0 | 21.4 | 5.47 | 25.5 | 3 |
| Tax 8F | LLFGYPV FV | 37.1 | 12.9 | 34.7 | 60.0 | 20.5 | 34.1 | 3 |
| Tax 5H | LLFGHPVYV | 36.2 | 13.1 | 36.2 | 56.1 | 8.23 | 14.7 | 3 |
| Tax 7H | LLFGYPHYV | 45.8 | 18.0 | 39.3 | 67.8 | 27.4 | 40.4 | 4 |
| Tax 4A | LLFAYPVYV | 72.5 | 5.28 | 7.3 | 118 | 18.9 | 16.0 | 3 |
| Tax 8A | LLFGYPVAV | 599 | 688 | 114.8 | 565 | 66.0 | 11.7 | 3 |
| Tax 4A8A | LLFAYPVAV | 194 | 207 | 106.6 | 1869 | 561 | 30.0 | 3 |
| Tax 5H8A | LLFGHPVAV | 2816 | 4292 | 152.4 | 2321 | 513 | 22.1 | 3 |

**Figure S1:**
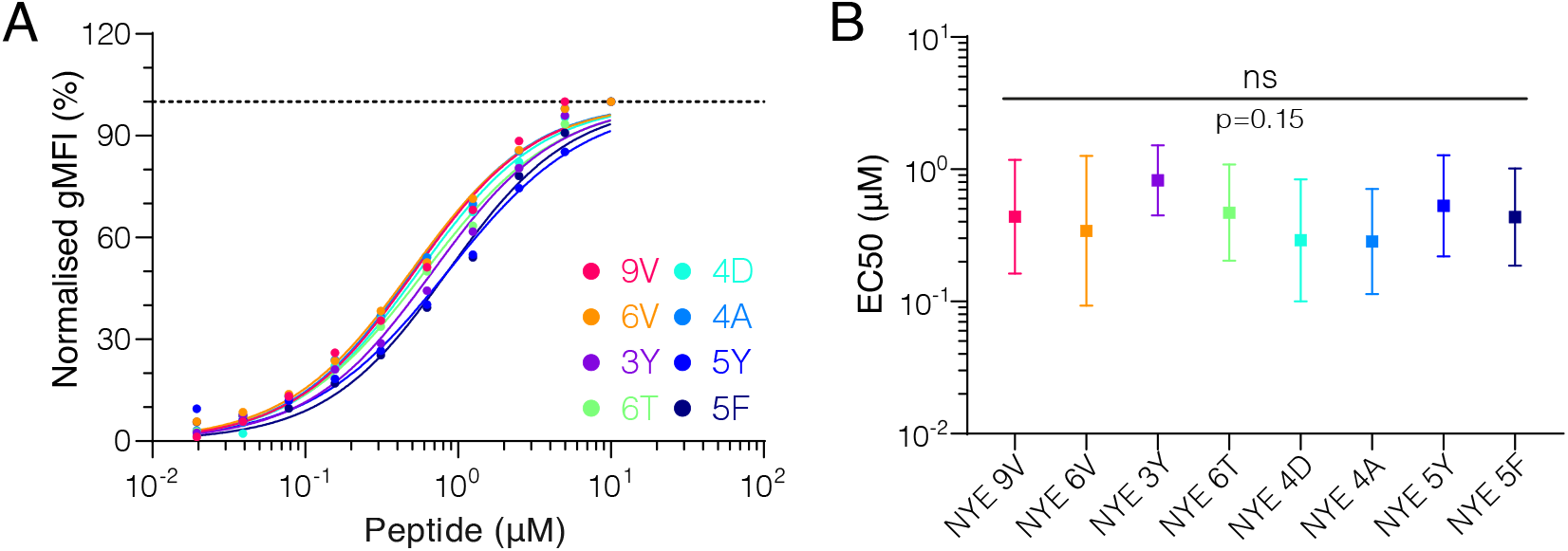
All NYE peptides load similarly on T2 cells. **(A)** Example of upregulation of HLA-A2 on T2 cells over different peptide pulsing concentrations. **(B)** Summary EC_50_s of loading on T2 cells. Shown are means with standard deviation. Pooled log-transformed data from 3 independent experiments were compared using a repeated-measure 1-way ANOVA with Geisser-Greenhouse correction.

**Figure S2:**
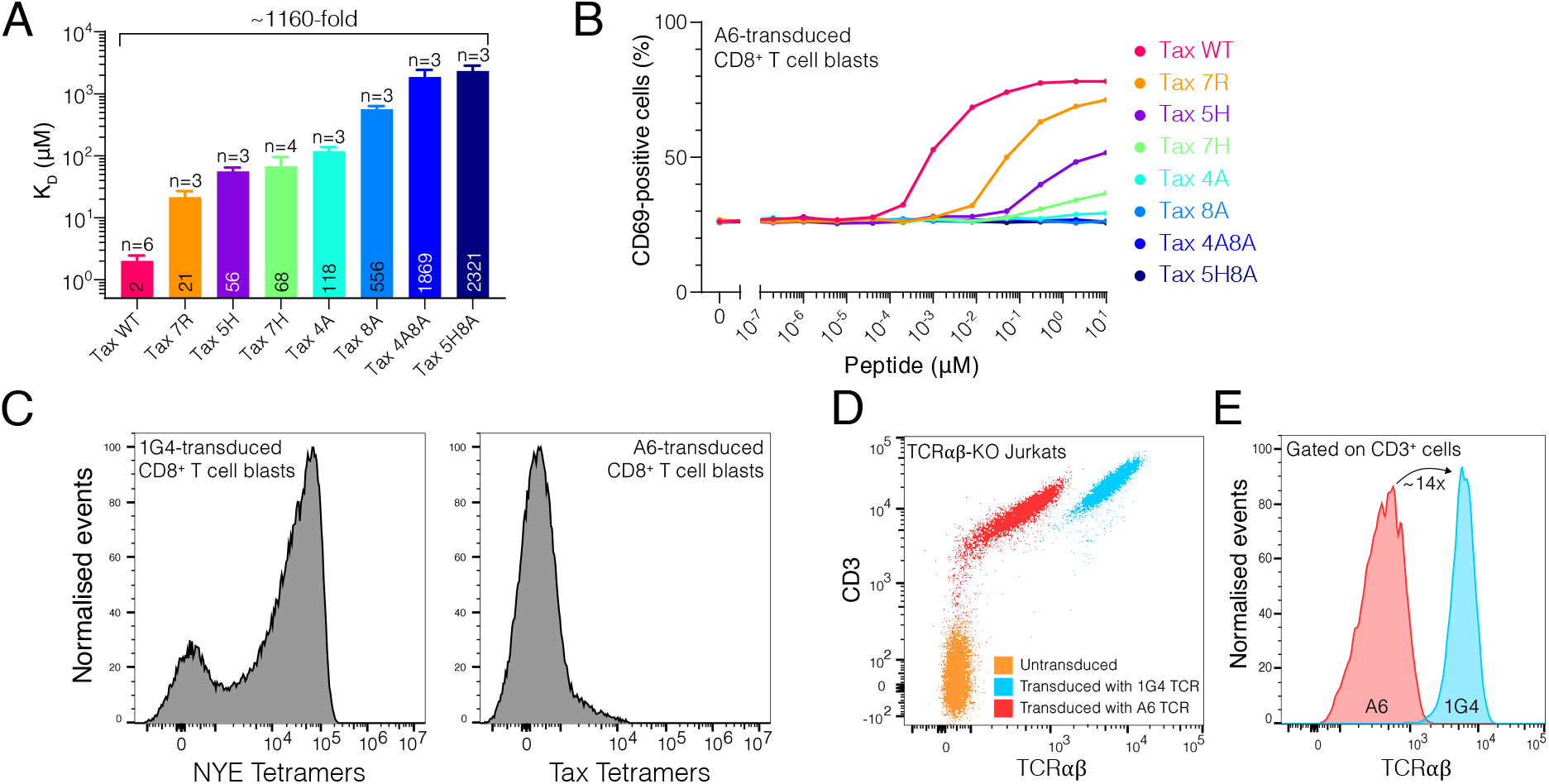
T cells transduced with A6 TCR have low expression and do not respond to ultra-low affinity pMHCs. **(A)** Affinity of the A6 TCR for the panel of Tax-related peptide, as measured by SPR. Shown are means with SD (the mean is also shown as a number wihtin the bar). **(B)** Activation of A6 expressing CD8^+^ T cell blasts by U87 APCs pulsed with the indicated concentration of Tax-related peptides. **(C)** Binding of the indicated pMHC tetramers to A6-versus 1G4-transduced CD8^+^ T cell blasts, as measured by flow cytometry. **(D)** Measurement by flow cytometry of cell surface expression of CD3 and TCRαβ by CD8^+^ T cell blasts following transduction with the indicated TCR into TCRαβ-KO Jurkats. **(E)** Relative expression of TCRαβ using the data in (E), gated on CD3^+^ cells.

**Figure S3:**
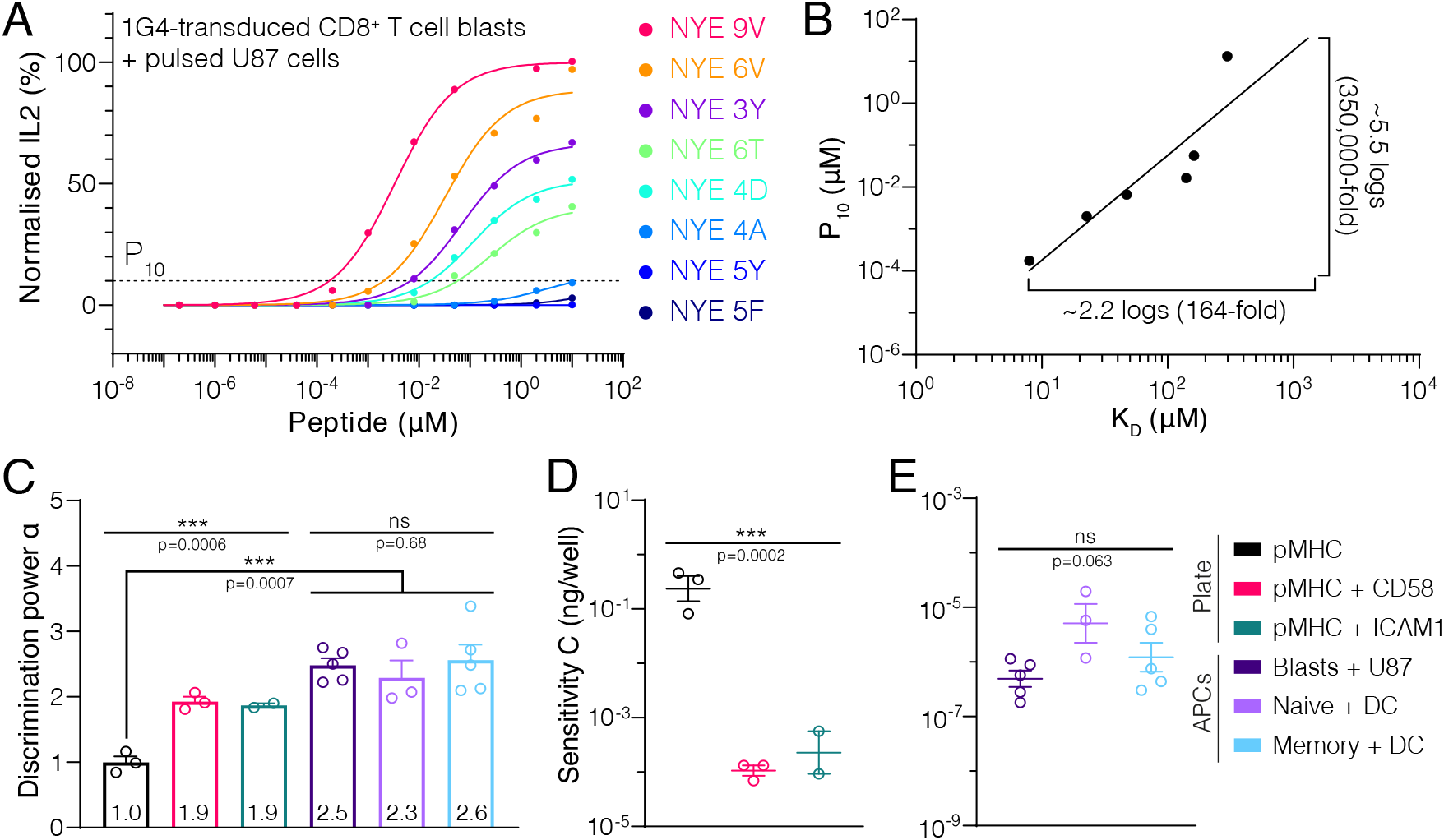
The low discrimination power is also observed for cytokine responses. **(A)** Dose-response for secretion of IL2 into the supernatant. Dotted line indicates 10 % activation threshold (*P*_10_). **(B)** A plot of potency obtained from **(A)** over affinity showing the power law fit. Lower affinity antigens that did not induce detectable IL-2 (NYE 5Y and 5F) are not included in the plot. **(C)** Discrimination power (α). Shown are means with SEM (value shown within bars). **(D,E)** Sensitivity measure (C, see main text) for stimulation with immobilised ligands **(D)** or APC co-culture **(E)**. Shown are means with SEM.

**Figure S4:**
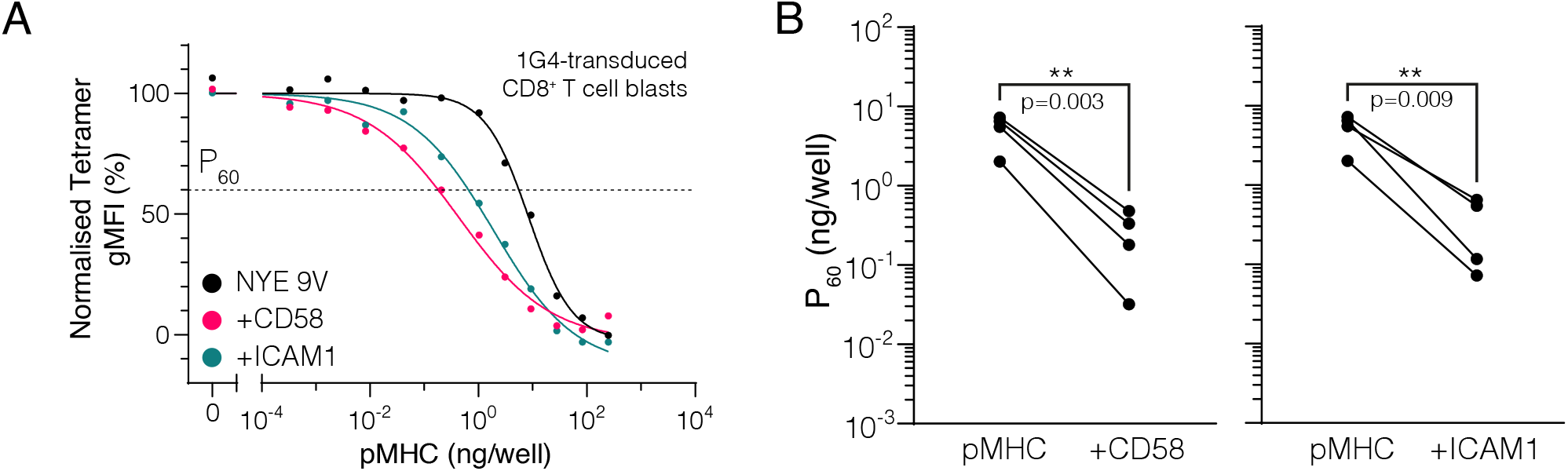
Engagement of CD2 or LFA-1 increases TCR downregulation. **(A)** Example dose-dependent for downregulation of TCR and its acceleration by engagement of CD2 by CD58 or LFA-1 by ICAM1. P_60_ is defined as the dose required to induce 40 % downregulation. Normalised to pMHC alone data. **(B)** Summary effect of CD2 or LFA-1 on TCR downregulation. Each dot represent an individual experiment. Statistics by repeated-measure 1-way ANOVA (with Geisser-Greenhouse correction) of log-transformed data (n=4) with Dunnett’s multiple comparison test.

**Figure S5:**
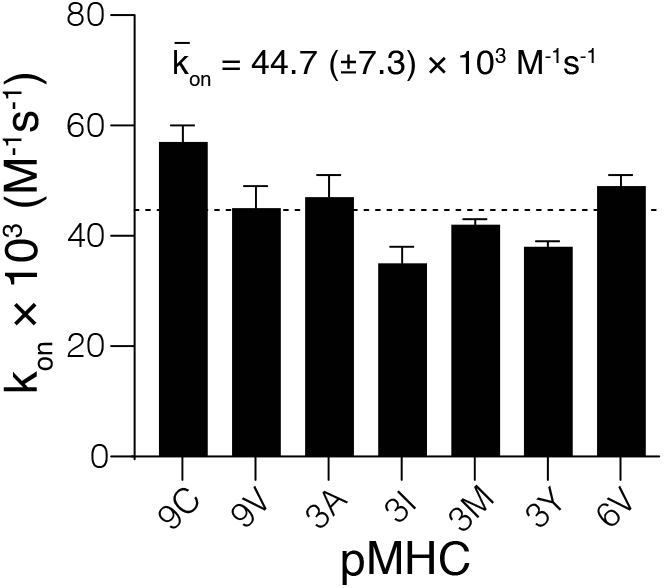
On-rates of 1G4 TCR with NYE peptides. Reported on-rates for 1G4 TCR with indicated NYE peptides as reported by (31). Shown are means with SEMs with 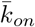 indicating the mean with SD. Lower affinity peptides (K_D_>100 *μ*M) were excluded.

**Figure S6:**
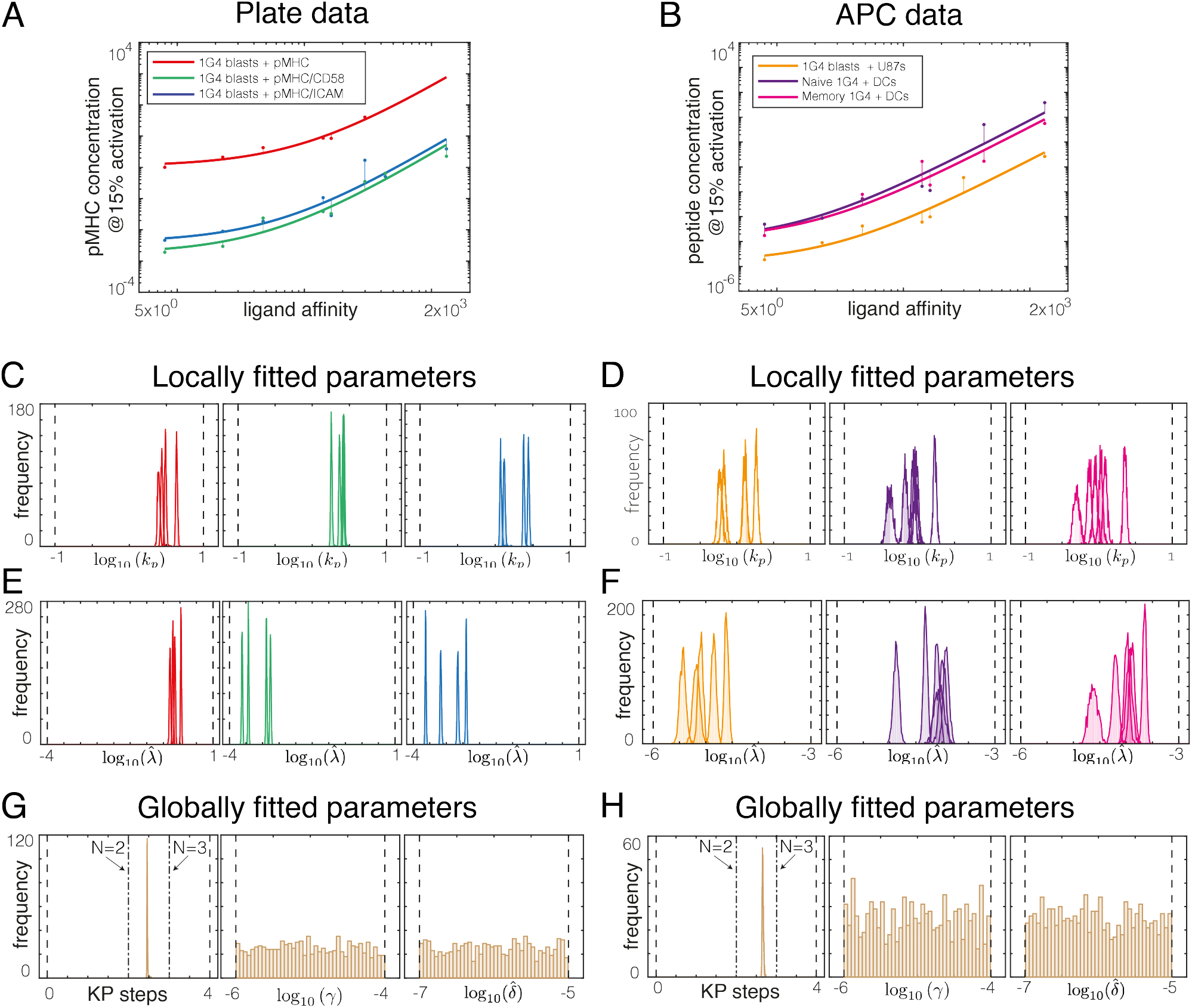
Direct fit of the kinetic proofreading model to potency data using the ABC-SMC method. **(A,B)** Examples of **(A)** plate and **(B)** APC potency data (dots) fitted with the KP model (lines). **(C-H)** The ABC-SMC method provides a distribution of all parameters that are consistent with the high quality fits presented in A and B. **(C-F)** Distribution of locally fitted parameters reveal that the proofreading rate (*k_p_*) and the sensitivity parameter 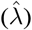 can be uniquely determined for each **(C, E)** plate experiment and **(D, F)** APC experiment. **(G-H)** Distribution of globally fitted parameters reveal that the number of KP steps (N) can be uniquely determined but that two additional parameter (*γ*, 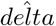) cannot be uniquely determined as they do not exhibit peaked distributions for **(G)** plate experiments and **(H)** APC experiments. See Supplementary Text for details.

**Figure S7:**
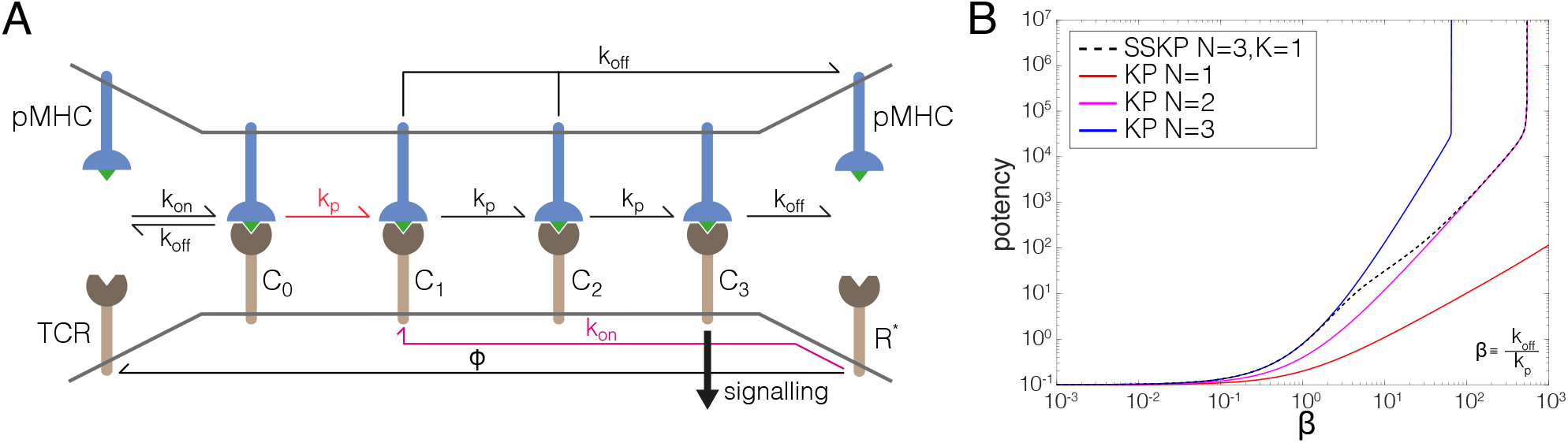
Kinetic proofreading with sustained signalling can explain apparent fractional steps. **(A)** Schematic of a 3-step kinetic proofreading model with sustained signalling in the 1^st^ step so that unbinding of pMHC after this step produces free TCR that sustains the 1^st^-step modification (state *R*\*). The binding of pMHC to this TCR can now short-circuit proofreading because only 2-steps are needed to reach the final signalling state (*C*_3_). The TCR sustains the signal when free for an average time of 1/*φ* and when *φ* becomes large this model reduces to the standard kinetic proofreading model. **(B)** Potency as a function of the *k*_off_/*k_p_* ratio (defined as *β*) for the standard kinetic proofreading model with *N* = 1, 2 or 3 steps (solid lines) or kinetic proofreading with sustained signalling with *N* = 3 steps and sustained signalling of the *K* = 1 step (dashed line).

## Supplementary Text

### Deriving the expression for potency: the kinetic proofreading model

The standard kinetic proofreading model (KP) for T cell receptor activation is as follows. A pMHC ligand *L* can bind with a T cell receptor *R* to create a complex *C*_0_ at a rate *k*_on_. In order for this complex to initiate an active T cell response it must undergo a series of biochemical modifications. These modifications are modelled by kinetic proofreading steps of which there are a total of *N*. We denote by *C_i_* a complex which is in the *i*-th KP step. A complex *C_i_* becomes a complex *C_i_*_+1_ with a progression rate *k_p_*, for 0 ≤ *i* ≤ *N* − 1. At any KP step the pMHC ligand can detach from the complex at rate *k*_off_. Let *L*(*t*), *R*(*t*), and *C_i_*(*t*) be the concentration of ligand, receptor and complex in the *i*-th KP step at time *t*, respectively. The system of ordinary differential equations that govern the temporal evolution of the concentrations is given by

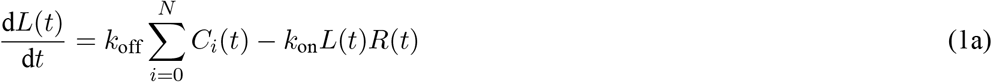

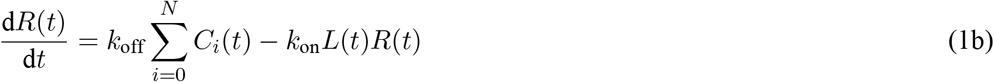

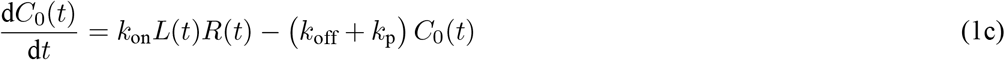

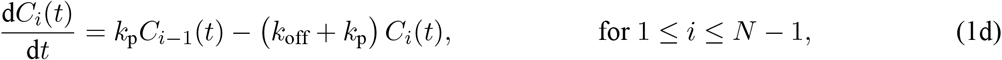

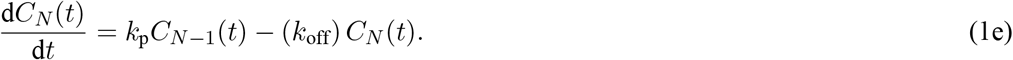

Let the initial number of pMHC ligands and T cell receptors be *L*_0_ and *R*_0_, respectively. We then define the total number of complexes at time *t* as 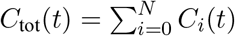, and note that we have two conservation equations, *L*_0_ = *L*(*t*) + *C*_tot_(*t*) and *R*_0_ = *R*(*t*) + *C*_tot_(*t*). Solving the steady state equations arising from setting the time derivatives in Eq. (1) to zero, and substituting in the conservation equations we find that

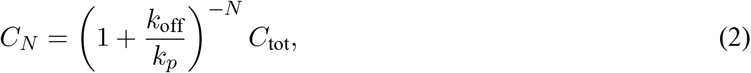

where

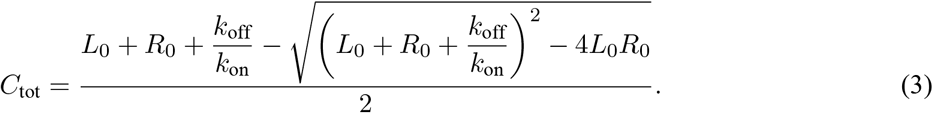

The expression in Eq. (2) determines the concentration of complex *C_N_*, which represents the strength of the activation signal, for a given number of ligands *L*_0_. To fit this model to the potency data seen in the main text we are interested in calculating the concentration of pMHC ligand required to initiate a T cell response given its binding properties. We first introduce a few convenient rescalings and redefinitions. We define *x* = *L*_0_/*R*_0_ to be the potency of ligand concentration relative to the total number of receptors and let *λ* = *C_N_* /*R*_0_ be a threshold parameter that dictates how much *C_N_* complex is needed to activate a T cell response relative to the total number of receptors. Thus Eq. (2) can be rewritten as

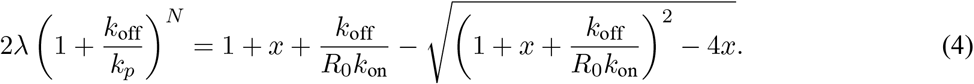

The experimental measurements of potency do not directly correspond to the potency *x* in our model as the exact number of ligand and receptor is unknown. Therefore we introduce a constant of proportionality *γ* into our model, such that *x* → *γx*. Similarly, the ratio *k*_off_/*k*_on_ is a measure of ligand affinity and is directly proportional to the experimental K_D_ values, thus we introduce a second constant of proportionality *δ* such that *k*_off_/(*R*_0_*k*_on_) → *δK_D_*, where we absorb the constant *R*_0_ into the new parameter. With these adjustments Equation (4) becomes

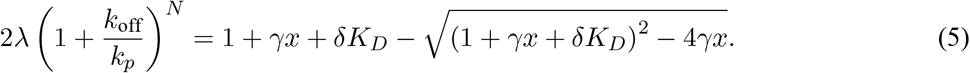

Upon rearranging Eq. (5) we find that

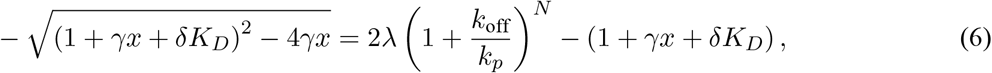

we then square^1^ both sides of Eq. (6) and find the following expression for the potency

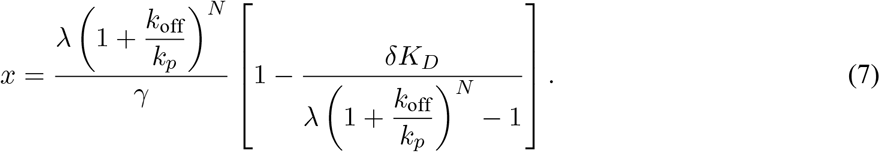

### ABC-SMC parameter estimation

Here we detail the Approximate Bayesian Computation-Sequential Monte Carlos algorithm used to determine the distribution of KP model parameters that fit the experimental data. Our KP model has five parameters, *N*, *k_p_*, *λ*, *γ* and *δ*. We fit the model parameters to the plate and the cell data separately. For both the plate and the cell data we fit *N*, *γ* and *δ* as a global parameter shared amongst all experimental runs. The parameters *k_p_* and *λ* are fitted locally for each run. We fit the potency equation to the experimental data in log space as such the log expression for potency, *ρ* (*N, k_p_*, 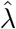, *γ*, 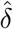),

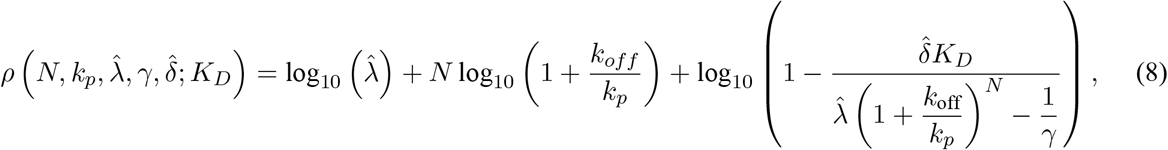

where 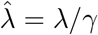 and 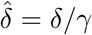. These rescalings ensure that the parameters are orthogonal and thus parameter space can be searched efficiently.

To perform ABC-SMC we first need to choose a prior distribution to initially sample the parameters. We chose uniform distributions that assume no prior knowledge about the system. However other than the parameter *N*, the parameters are uniform in log space. This allows for efficient search through parameter space over many orders of magnitude. The priors for the plate data are as follows

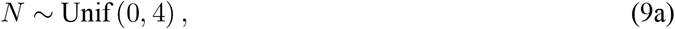

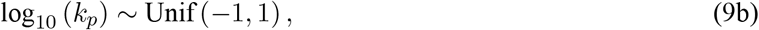

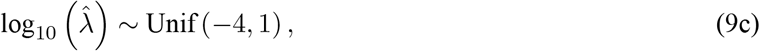

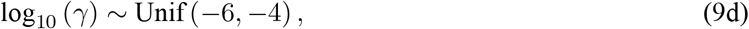

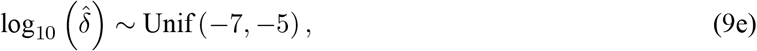

where the priors for the cell data are the same other than for 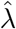 where 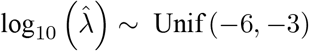.

Recall that we fit the parameters *N*, *γ*, and 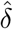 globally and 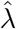 and *k_p_* are fitted locally. For the plate data this results in 27 parameters to fit whilst for the cell data there are 37 parameters. Let 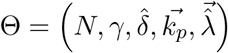 be the vector of parameters to fit such that the *i*-th entry of the vectors 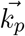 and 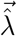 correspond to the local parameters for the *i*-th experiment. Then let 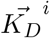 be the vector of experimentally measured K_D_ values, and 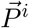 be the vector of potency measurements for the *i*-th experiment. These vectors differ in length and so we denote by *d_i_* the number of data points in the *i*-th experiment. We measure the similarity between the KP model and the experimental results via the following distance function

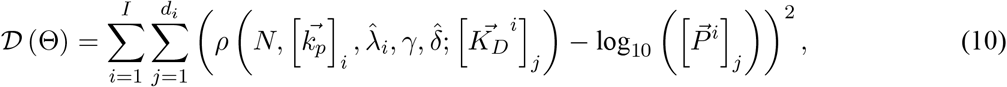

where *I* denotes the total number of experiments, *I* = 12 and *I* = 17 for the plate and cell data, respectively.

To perform a randomised search through the parameter space we employed the following Metropolis-Hastings algorithm. We sample an initial parameter set Θ_0_ from the prior distributions detailed above. Let Θ_curr_ denote the current set of parameters which initially is Θ_0_. A candidate set of parameters, Θ_cand_ is found by adding a random perturbation to Θ_curr_. The perturbation is achieved by adding a uniform random shift to each parameter in Θ_curr_ independently.The range of the uniform random shift is [−0.005, 0.005] multiplied by the width of the prior. For example we perturb the *N* parameter by adding a random uniform shift in the interval [−0.02, 0.02]. If the parameter falls outside the bounds in the prior distribution it is reflected symmetrically back within the bounds. We then have to decide whether to accept or reject the candidate set of parameters. If 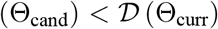 then we accept the parameters as they share a greater similarity with the experimental data and set Θ_curr_ = Θ_cand_. Otherwise we only accept the candidate parameters with probability 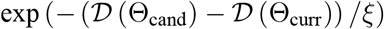, where *ξ* is a parameter that controls how likely accepting a set of parameters with a higher distance function is. The value of *ξ* is reduced as the algorithm gets closer to a set of parameters that minimises the distance function. Initially *ξ* = 10 but is subsequently reduced to {1, 0.1, 0.01, 0.005, 0.001} when the distance function of the candidate set of parameters first reaches {50, 30, 20, 18, 17.5} for the plate data and {100, 75, 50, 40, 35} for the cell data. The algorithm continues until it reaches a final set of parameters that has a distance less than 11.08 or 39.2 for the plate and cell data, respectively. For both the plate and cell data we performed this algorithm 1000 times to capture the distribution of parameter values that fit the experimental data.

### Kinetic proofreading with sustained signalling

The sustained signaling kinetic proofreading model (SSKP) is a generalization of the standard kinetic proofreading model (KP), where one of the proofreading steps is difficult to revert and upon ligand dissociation there is a temporary window where a ligand can rebind to a sustained receptor and bypass some of the initial KP steps. To be precise, let there be *N* KP steps where the *K*-th step out of *N* is sustained. As before *C_i_* denotes a complex in the *i*-th KP step, and *L*(*t*) and *R*(*t*) denote the concentration of pMHC ligand and TCR, respectively at time *t*. In addition, we introduce *R*\*(*t*) as the concentration of unbound receptor that sustains the *K*-th modification. A ligand can bind with this receptor at rate *k*_on_ to form complex *C_K_*, rather than *C*_0_. The system of ordinary differential equations that governs the temporal evolution of the concentrations is given by

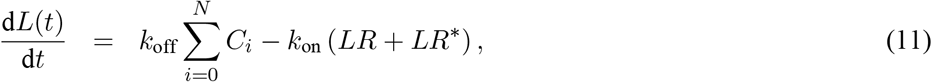

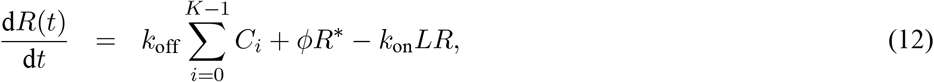

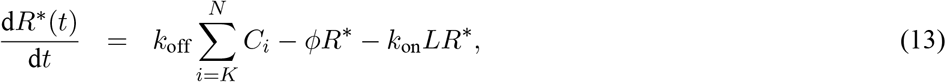

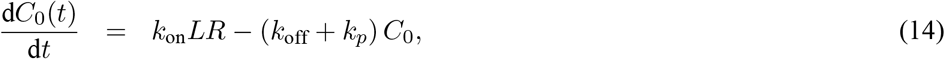

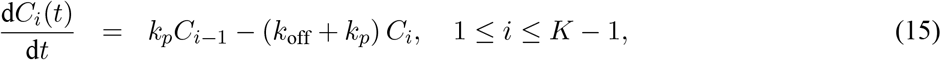

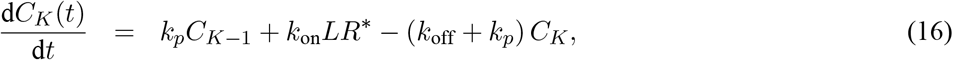

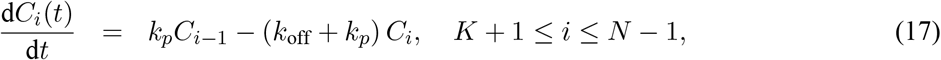

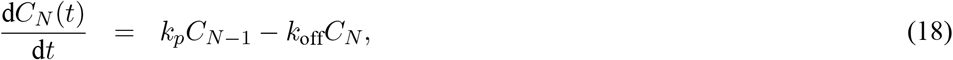

The initial number of pMHC ligands and TCRs is given by *L*_0_ and *R*_0_, respectively. As before 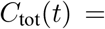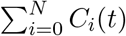 denotes the total concentration of complexes. By setting the time derivatives to zero in the above system of ODEs we calculate the equilibrium concentration of *C_N_* to be

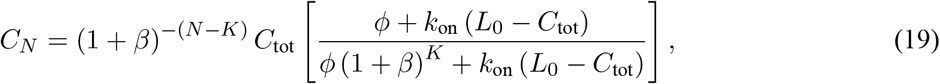

where *β* = *k*_off_/*k_p_* and

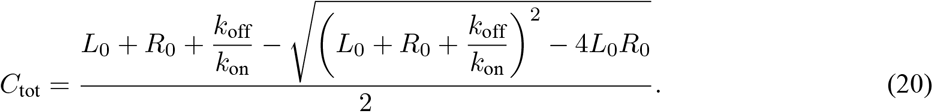

Firstly, we note that when *φ* → ∞ Equation (19) becomes (1 + *β*)^−*N*^ *C*_tot_ which is the standard KP model result, as to be expected. If we set *φ* = 0 we get *C_N_* = (1 + *β*)^−(*N*−*K*)^ *C*_tot_, this is the result for standard KP with *N* − *K* steps. Thus, for *φ* nonzero and non-infinite we expect to see an effective number of KP steps in-between *N* and *N* − *K*.

Unlike the standard KP model there is no explicit formula for extracting a potency plot, the required ligand for activation as a function of the dissociation rate. Instead, the potency plot in Figure S7B was created by solving *C_N_* = *λ* from Equation (19) numerically to extract *L*_0_ the minimum ligand concentration that guarantees the signaling complex *C_N_* reaches a threshold *λ*.

## Footnotes

1 Squaring both sides will not introduce a false solution so long as 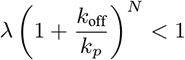.

