## Supplementary Information for "T cells exhibit unexpectedly low discriminatory power and can respond to ultra-low affinity peptide-MHC ligands"

### Supplementary Text

#### Deriving the expression for potency: the kinetic proofreading model

The standard kinetic proofreading model (KP) for T cell receptor activation is as follows. A pMHC ligand $L$ can bind with a T cell receptor $R$ to create a complex $C_0$ at a rate $k_{\text{on}}$. In order for this complex to initiate an active T cell response it must undergo a series of biochemical modifications. These modifications are modelled by kinetic proofreading steps of which there are a total of $N$. We denote by $C_i$ a complex which is in the $i$-th KP step. A complex $C_i$ becomes a complex $C_{i+1}$ with a progression rate $k_p$, for $0 \leq i \leq N-1$. At any KP step the pMHC ligand can detach from the complex at rate $k_{\text{off}}$. Let $L(t)$, $R(t)$, and $C_i(t)$ be the concentration of ligand, receptor and complex in the $i$-th KP step at time $t$, respectively. The system of ordinary differential equations that govern the temporal evolution of the concentrations is given by

$$\frac{\mathrm{d}L(t)}{\mathrm{d}t} = k_{\text{off}} \sum_{i=0}^{N} C_i(t) - k_{\text{on}} L(t) R(t) \tag{1a}$$

$$\frac{\mathrm{d}R(t)}{\mathrm{d}t} = k_{\text{off}} \sum_{i=0}^{N} C_i(t) - k_{\text{on}} L(t) R(t) \tag{1b}$$

$$\frac{\mathrm{d}C_0(t)}{\mathrm{d}t} = k_{\text{on}} L(t) R(t) - \left(k_{\text{off}} + k_{\text{p}}\right) C_0(t) \tag{1c}$$

$$\frac{\mathrm{d}C_i(t)}{\mathrm{d}t} = k_{\text{p}} C_{i-1}(t) - \left(k_{\text{off}} + k_{\text{p}}\right) C_i(t), \qquad \text{for } 1 \leq i \leq N-1, \tag{1d}$$

$$\frac{\mathrm{d}C_N(t)}{\mathrm{d}t} = k_{\text{p}} C_{N-1}(t) - \left(k_{\text{off}}\right) C_N(t). \tag{1e}$$

Let the initial number of pMHC ligands and T cell receptors be $L_0$ and $R_0$, respectively. We then define the total number of complexes at time $t$ as $C_{\text{tot}}(t) = \sum_{i=0}^{N} C_i(t)$, and note that we have two conservation equations, $L_0 = L(t) + C_{\text{tot}}(t)$ and $R_0 = R(t) + C_{\text{tot}}(t)$. Solving the steady state equations arising from setting the time derivatives in Eq. (1) to zero, and substituting in the conservation equations we find that

$$C_N = \left(1 + \frac{k_{\text{off}}}{k_p}\right)^{-N} C_{\text{tot}}, \tag{2}$$

where

$$C_{\text{tot}} = \frac{L_0 + R_0 + \dfrac{k_{\text{off}}}{k_{\text{on}}} - \sqrt{\left(L_0 + R_0 + \dfrac{k_{\text{off}}}{k_{\text{on}}}\right)^2 - 4L_0 R_0}}{2}. \tag{3}$$

The expression in Eq. (2) determines the concentration of complex $C_N$, which represents the strength of the activation signal, for a given number of ligands $L_0$. To fit this model to the potency data seen in the main text we are interested in calculating the concentration of pMHC ligand required to initiate a T cell response given its binding properties. We first introduce a few convenient rescalings and redefinitions. We define $x = L_0/R_0$ to be the potency of ligand concentration relative to the total number of receptors and let $\lambda = C_N/R_0$ be a threshold parameter that dictates how much $C_N$ complex is needed to activate a T cell response relative to the total number of receptors. Thus Eq. (2) can be rewritten as

$$2\lambda\left(1+\frac{k_{\text{off}}}{k_p}\right)^N = 1+x+\frac{k_{\text{off}}}{R_0 k_{\text{on}}} - \sqrt{\left(1+x+\frac{k_{\text{off}}}{R_0 k_{\text{on}}}\right)^2 - 4x}. \tag{4}$$

The experimental measurements of potency do not directly correspond to the potency $x$ in our model as the exact number of ligand and receptor is unknown. Therefore we introduce a constant of proportionality $\gamma$ into our model, such that $x \to \gamma x$. Similarly, the ratio $k_{\text{off}}/k_{\text{on}}$ is a measure of ligand affinity and is directly proportional to the experimental $K_D$ values, thus we introduce a second constant of proportionality $\delta$ such that $k_{\text{off}}/(R_0 k_{\text{on}}) \to \delta K_D$, where we absorb the constant $R_0$ into the new parameter. With these adjustments Equation (4) becomes

$$2\lambda\left(1+\frac{k_{\text{off}}}{k_p}\right)^N = 1+\gamma x+\delta K_D - \sqrt{(1+\gamma x+\delta K_D)^2 - 4\gamma x}. \tag{5}$$

Upon rearranging Eq. (5) we find that

$$-\sqrt{(1+\gamma x+\delta K_D)^2 - 4\gamma x} = 2\lambda\left(1+\frac{k_{\text{off}}}{k_p}\right)^N - (1+\gamma x+\delta K_D)\,, \tag{6}$$

we then square[1] both sides of Eq. (6) and find the following expression for the potency

$$x = \frac{\lambda\left(1+\frac{k_{\text{off}}}{k_p}\right)^N}{\gamma}\left[1 - \frac{\delta K_D}{\lambda\left(1+\frac{k_{\text{off}}}{k_p}\right)^N - 1}\right]. \tag{7}$$

#### ABC-SMC parameter estimation

Here we detail the Approximate Bayesian Computation-Sequential Monte Carlos algorithm used to determine the distribution of KP model parameters that fit the experimental data. Our KP model has five parameters, $N$, $k_p$, $\lambda$, $\gamma$ and $\delta$. We fit the model parameters to the plate and the cell data separately. For both the plate and the cell data we fit $N$, $\gamma$ and $\delta$ as a global parameter shared amongst all experimental runs. The parameters $k_p$ and $\lambda$ are fitted locally for each run. We fit the potency equation to the experimental data in log space as such the log expression for potency, $\rho\left(N, k_p, \hat{\lambda}, \gamma, \hat{\delta}\right)$,

$$\rho\left(N, k_p, \hat{\lambda}, \gamma, \hat{\delta}; K_D\right) = \log_{10}\left(\hat{\lambda}\right) + N\log_{10}\left(1+\frac{k_{off}}{k_p}\right) + \log_{10}\left(1 - \frac{\hat{\delta}K_D}{\hat{\lambda}\left(1+\frac{k_{\text{off}}}{k_p}\right)^N - \frac{1}{\gamma}}\right), \tag{8}$$

where $\hat{\lambda} = \lambda/\gamma$ and $\hat{\delta} = \delta/\gamma$. These rescalings ensure that the parameters are orthogonal and thus parameter space can be searched efficiently.

To perform ABC-SMC we first need to choose a prior distribution to initially sample the parameters. We chose uniform distributions that assume no prior knowledge about the system. However other than the parameter $N$,

[1] Squaring both sides will not introduce a false solution so long as $\lambda\left(1+\frac{k_{\text{off}}}{k_p}\right)^N < 1$.

the parameters are uniform in log space. This allows for efficient search through parameter space over many orders of magnitude. The priors for the plate data are as follows

$$N \sim \text{Unif}\,(0,4)\,, \tag{9a}$$
$$\log_{10}(k_p) \sim \text{Unif}\,(-1,1)\,, \tag{9b}$$
$$\log_{10}\left(\hat{\lambda}\right) \sim \text{Unif}\,(-4,1)\,, \tag{9c}$$
$$\log_{10}(\gamma) \sim \text{Unif}\,(-6,-4)\,, \tag{9d}$$
$$\log_{10}\left(\hat{\delta}\right) \sim \text{Unif}\,(-7,-5)\,, \tag{9e}$$

where the priors for the cell data are the same other than for $\hat{\lambda}$ where $\log_{10}\left(\hat{\lambda}\right) \sim \text{Unif}\,(-6,-3)$.

Recall that we fit the parameters $N$, $\gamma$, and $\hat{\delta}$ globally and $\hat{\lambda}$ and $k_p$ are fitted locally. For the plate data this results in 27 parameters to fit whilst for the cell data there are 37 parameters. Let $\Theta = \left(N, \gamma, \hat{\delta}, \vec{k_p}, \vec{\hat{\lambda}}\right)$ be the vector of parameters to fit such that the $i$-th entry of the vectors $\vec{k_p}$ and $\vec{\hat{\lambda}}$ correspond to the local parameters for the $i$-th experiment. Then let $\vec{K_D}^{i}$ be the vector of experimentally measured $K_D$ values, and $\vec{P}^{i}$ be the vector of potency measurements for the $i$-th experiment. These vectors differ in length and so we denote by $d_i$ the number of data points in the $i$-th experiment. We measure the similarity between the KP model and the experimental results via the following distance function

$$\mathcal{D}\,(\Theta) = \sum_{i=1}^{I}\sum_{j=1}^{d_i}\left(\rho\left(N, \left[\vec{k_p}\right]_i, \hat{\lambda}_i, \gamma, \hat{\delta}; \left[\vec{K_D}^{i}\right]_j\right) - \log_{10}\left(\left[\vec{P}^{i}\right]_j\right)\right)^2, \tag{10}$$

where $I$ denotes the total number of experiments, $I = 12$ and $I = 17$ for the plate and cell data, respectively.

To perform a randomised search through the parameter space we employed the following Metropolis-Hastings algorithm. We sample an initial parameter set $\Theta_0$ from the prior distributions detailed above. Let $\Theta_{\text{curr}}$ denote the current set of parameters which initially is $\Theta_0$. A candidate set of parameters, $\Theta_{\text{cand}}$ is found by adding a random perturbation to $\Theta_{\text{curr}}$. The perturbation is achieved by adding a uniform random shift to each parameter in $\Theta_{\text{curr}}$ independently.The range of the uniform random shift is $[-0.005, 0.005]$ multiplied by the width of the prior. For example we perturb the $N$ parameter by adding a random uniform shift in the interval $[-0.02, 0.02]$. If the parameter falls outside the bounds in the prior distribution it is reflected symmetrically back within the bounds. We then have to decide whether to accept or reject the candidate set of parameters. If $\mathcal{D}\,(\Theta_{\text{cand}}) < \mathcal{D}\,(\Theta_{\text{curr}})$ then we accept the parameters as they share a greater similarity with the experimental data and set $\Theta_{\text{curr}} = \Theta_{\text{cand}}$. Otherwise we only accept the candidate parameters with probability $\exp\left(-\left(\mathcal{D}\,(\Theta_{\text{cand}}) - \mathcal{D}\,(\Theta_{\text{curr}})\right)/\xi\right)$, where $\xi$ is a parameter that controls how likely accepting a set of parameters with a higher distance function is. The value of $\xi$ is reduced as the algorithm gets closer to a set of parameters that minimises the distance function. Initially $\xi = 10$ but is subsequently reduced to $\{1, 0.1, 0.01, 0.005, 0.001\}$ when the distance function of the candidate set of parameters first reaches $\{50, 30, 20, 18, 17.5\}$ for the plate data and $\{100, 75, 50, 40, 35\}$ for the cell data. The algorithm continues until it reaches a final set of parameters that has a distance less than 11.08 or 39.2 for the plate and cell data, respectively. For both the plate and cell data we performed this algorithm 1000 times to capture the distribution of parameter values that fit the experimental data.

#### Kinetic proofreading with sustained signalling

The sustained signaling kinetic proofreading model (SSKP) is a generalization of the standard kinetic proofreading model (KP), where one of the proofreading steps is difficult to revert and upon ligand dissociation there is a temporary window where a ligand can rebind to a sustained receptor and bypass some of the initial KP steps. To be precise, let there be $N$ KP steps where the $K$-th step out of $N$ is sustained. As before $C_i$ denotes a complex in the $i$-th KP step, and $L(t)$ and $R(t)$ denote the concentration of pMHC ligand and TCR, respectively at time $t$. In addition, we introduce $R^*(t)$ as the concentration of unbound receptor that sustains the $K$-th modification. A ligand can bind with this receptor at rate $k_{\text{on}}$ to form complex $C_K$, rather than $C_0$. The system of ordinary differential equations that governs the temporal evolution of the concentrations is given by

$$\frac{\mathrm{d}L(t)}{\mathrm{d}t} = k_{\text{off}} \sum_{i=0}^{N} C_i - k_{\text{on}} \left( LR + LR^* \right), \tag{11}$$

$$\frac{\mathrm{d}R(t)}{\mathrm{d}t} = k_{\text{off}} \sum_{i=0}^{K-1} C_i + \phi R^* - k_{\text{on}} LR, \tag{12}$$

$$\frac{\mathrm{d}R^*(t)}{\mathrm{d}t} = k_{\text{off}} \sum_{i=K}^{N} C_i - \phi R^* - k_{\text{on}} LR^*, \tag{13}$$

$$\frac{\mathrm{d}C_0(t)}{\mathrm{d}t} = k_{\text{on}} LR - \left( k_{\text{off}} + k_p \right) C_0, \tag{14}$$

$$\frac{\mathrm{d}C_i(t)}{\mathrm{d}t} = k_p C_{i-1} - \left( k_{\text{off}} + k_p \right) C_i, \quad 1 \leq i \leq K-1, \tag{15}$$

$$\frac{\mathrm{d}C_K(t)}{\mathrm{d}t} = k_p C_{K-1} + k_{\text{on}} LR^* - \left( k_{\text{off}} + k_p \right) C_K, \tag{16}$$

$$\frac{\mathrm{d}C_i(t)}{\mathrm{d}t} = k_p C_{i-1} - \left( k_{\text{off}} + k_p \right) C_i, \quad K+1 \leq i \leq N-1, \tag{17}$$

$$\frac{\mathrm{d}C_N(t)}{\mathrm{d}t} = k_p C_{N-1} - k_{\text{off}} C_N, \tag{18}$$

The initial number of pMHC ligands and TCRs is given by $L_0$ and $R_0$, respectively. As before $C_{\text{tot}}(t) = \sum_{i=0}^{N} C_i(t)$ denotes the total concentration of complexes. By setting the time derivatives to zero in the above system of ODEs we calculate the equilibrium concentration of $C_N$ to be

$$C_N = (1+\beta)^{-(N-K)} C_{\text{tot}} \left[ \frac{\phi + k_{\text{on}} \left( L_0 - C_{\text{tot}} \right)}{\phi \left( 1+\beta \right)^K + k_{\text{on}} \left( L_0 - C_{\text{tot}} \right)} \right], \tag{19}$$

where $\beta = k_{\text{off}}/k_p$ and

$$C_{\text{tot}} = \frac{L_0 + R_0 + \dfrac{k_{\text{off}}}{k_{\text{on}}} - \sqrt{\left( L_0 + R_0 + \dfrac{k_{\text{off}}}{k_{\text{on}}} \right)^2 - 4 L_0 R_0}}{2}. \tag{20}$$

Firstly, we note that when $\phi \to \infty$ Equation (19) becomes $(1+\beta)^{-N} C_{\text{tot}}$ which is the standard KP model result, as to be expected. If we set $\phi = 0$ we get $C_N = (1+\beta)^{-(N-K)} C_{\text{tot}}$, this is the result for standard KP with $N-K$ steps. Thus, for $\phi$ nonzero and non-infinite we expect to see an effective number of KP steps in-between $N$ and $N-K$.

Unlike the standard KP model there is no explicit formula for extracting a potency plot, the required ligand for activation as a function of the dissociation rate. Instead, the potency plot in Figure S7B was created by solving $C_N = \lambda$ from Equation (19) numerically to extract $L_0$ the minimum ligand concentration that guarantees the signaling complex $C_N$ reaches a threshold $\lambda$.
